# Systematic discovery of endogenous human ribonucleoprotein complexes

**DOI:** 10.1101/480061

**Authors:** Anna L. Mallam, Wisath Sae-Lee, Jeffrey M. Schaub, Fan Tu, Anna Battenhouse, Yu Jin Jang, Jonghwan Kim, John B. Wallingford, Ilya J. Finkelstein, Edward M. Marcotte, Kevin Drew

## Abstract

RNA-binding proteins (RBPs) play essential roles in biology and are frequently associated with human disease. While recent studies have systematically identified individual RBPs, their higher order assembly into <u>R</u>ibo<u>n</u>ucleo<u>p</u>rotein (RNP) complexes has not been systematically investigated. Here, we describe a proteomics method for systematic identification of RNP complexes in human cells. We identify 1,428 protein complexes that associate with RNA, indicating that over 20% of known human protein complexes contain RNA. To explore the role of RNA in the assembly of each complex, we identify complexes that dissociate, change composition, or form stable protein-only complexes in the absence of RNA. Importantly, these data also provide specific novel insights into the function of well-studied protein complexes not previously known to associate with RNA, including replication factor C (RFC) and cytokinetic centralspindlin complex. Finally, we use our method to systematically identify cell-type specific RNA-associated proteins in mouse embryonic stem cells. We distribute these data as a resource, rna.MAP (rna.proteincomplexes.org) which provides a comprehensive dataset for the study of RNA-associated protein complexes. Our system thus provides a novel methodology for further explorations across human tissues and disease states, as well as throughout all domains of life.

**Summary:** An exploration of human protein complexes in the presence and absence of RNA reveals endogenous ribonucleoprotein complexes

## Introduction

RNA-binding proteins (RBPs) play essential roles in diverse biological processes, and in most cases act within higher-order multi-protein complexes called <u>R</u>ibo<u>n</u>ucleo<u>p</u>rotein (RNP) complexes (Castello et al., 2013; Gerstberger et al., 2014; Hentze et al., 2018). Understanding RNPs is of particular importance due to their indispensable role in many essential cellular functions, such as mRNA splicing (spliceosome) (Wahl et al., 2009), translation (ribosome) (Ramakrishnan, 2002), silencing (RISC) (Kawamata and Tomari, 2010), and degradation (exosome) (Houseley et al., 2006). Moreover, RNPs also play more specific roles in, for example, mRNA transport and localization in developing embryos and mature neurons (Holt and Bullock, 2009; Sahoo et al., 2018) and assembly phase separated organelles (Mittag and Parker, 2018). Further, RNPs are strongly implicated in human diseases including amyotrophic lateral sclerosis (ALS) (Scotter et al., 2015), spinocerebellar ataxia (Yue et al., 2001), and autism (Voineagu et al., 2011). Accordingly, substantial recent effort has been focused on systematic identification of RNA-associated proteins (Baltz et al., 2012; Bao et al., 2018; Brannan et al., 2016; Castello et al., 2012, 2016; He et al., 2016; Huang et al., 2018; Queiroz et al., 2019; Treiber et al., 2017; Trendel et al., 2019).

Strikingly, however, we still lack any systematic characterization of the assembly of individual RNA-associated proteins into the higher order RNP complexes in which so many function, leaving a crucial gap in our knowledge. A worldwide effort is currently underway to systematically identify multi-protein complexes using high throughput mass spectrometry techniques (Hein et al., 2015; Huttlin et al., 2015), but none of these techniques identify an RNA component within the complexes. We therefore set out to develop a method for systematic identification of RNA-associated higher-order multi-protein complexes that requires no genetic manipulation (i.e. tag-free) and would be easily adaptable to diverse cell types.

Here, we present ‘<u>dif</u>ferential <u>frac</u>tionation for interaction analysis’ (DIF-FRAC), which measures the sensitivity of protein complexes to a given treatment (ex. RNase A) using native size-exclusion chromatography followed by mass spectrometry. DIF-FRAC is based on a high throughput co-fractionation mass spectrometry (CF-MS) approach that we developed and applied to a diverse set of tissues and cells types en route to generating human and metazoan protein complex maps (Drew et al., 2017; Havugimana et al., 2012; Wan et al., 2015). DIF-FRAC builds upon CF-MS by comparing chromatographic separations of cellular lysate under control and RNA degrading conditions (Figure 1A). A statistical framework is then applied to discover RNP complexes by identifying concurrent shifts of known protein complex subunits upon RNA degradation (Figure 1A).

**Figure 1:**
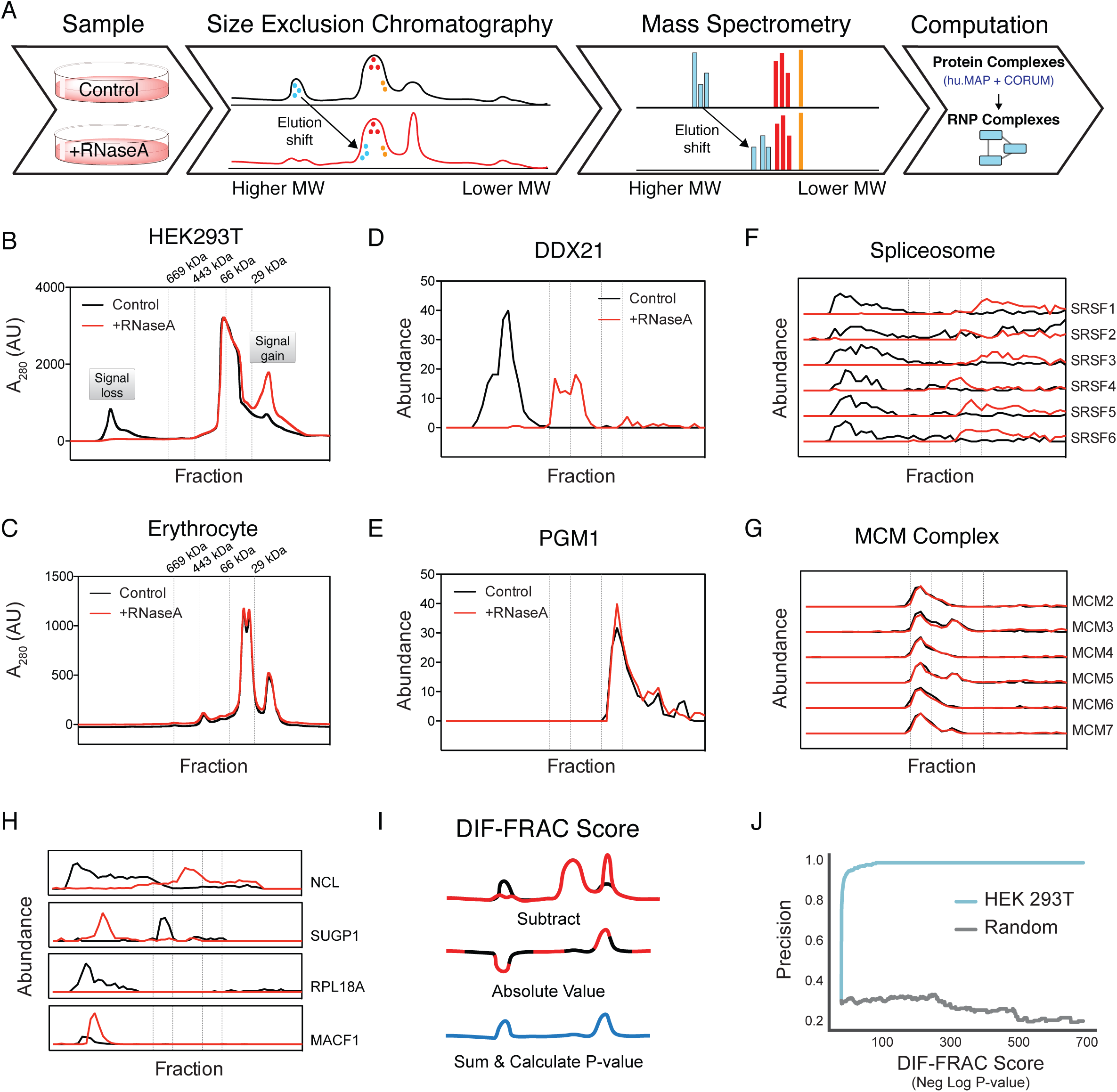
<u>Dif</u>ferential <u>frac</u>tionation (DIF-FRAC) identifies RNP complexes. (A) The DIF-FRAC workflow requires two equivalent cell culture lysates for a control and an RNase A treated sample. Lysate is separated into fractions by size exclusion chromatography (SEC), and proteins in each fraction are identified by mass spectrometry to determine individual protein elution profiles proteome-wide for each condition. An elution shift of a protein upon RNase A treatment is indicative of an RNA-protein association. Elution shifts are cross-referenced with known protein complexes to identify RNP complexes. (B) Separations of HEK293T lysate under control (black) and RNase A treated (red) conditions monitored by bulk SEC chromatography absorbance profiles at A_280_ show loss of high molecular weight signal upon treatment. (C) Negative control separations of erythrocyte lysate under control (black) and RNase A treated (red) conditions monitored by bulk SEC chromatography absorbance profiles at A_280_ show no change in absorbance signal. (D) RNA binding protein elution profile for positive control nucleolar RNA helicase 2 (DDX21) (Abundance = count of unique peptide spectral matches). The elution profile shows sensitivity to RNase A treatment. (E) Elution profile for negative control phosphoglucomutase (PGM1) is not sensitive to RNase A treatment. (F) Elution profiles for subunits of the spliceosome RNP complex (i.e. positive control) show co-elution of complex in control and a shift in elution upon RNase A treatment. (G) Elution profile for the non-RNA-associated MCM complex (i.e. negative control) shows no detectable elution shift. (H) Example traces of four known RNA-associated proteins exhibiting different behaviors of elution profile changes upon RNase A treatment. NCL shows a loss in molecular weight while SUGP1 shows an increase in molecular weight. RPL18A shows a decrease in observed abundance while MACF1 shows an increase in observed abundance. In (B)-(H) dashed lines correspond to the elution volumes of molecular weight standards thyroglobulin (M_r_ = 669 kDa), apoferritin (M_r_ = 443 kDa), albumin (M_r_ = 66 kDa), and carbonic anhydrase (M_r_ = 29 kDa). Molecular weight labels on subsequent plots are removed for clarity. (I) A DIF-FRAC score is calculated for each protein from the absolute value of the difference of the elution profiles between control and RNase A treated samples, and then summed. A P-value is then calculated from a background distribution of DIF-FRAC scores. See also Figure S3A. (J) DIF-FRAC P-value calculated on HEK 293T data shows strong ability to discriminate known RNA-associated proteins from other proteins. See also Figure S3B.

Analysis of DIF-FRAC data answers important questions as to the role of RNA plays in macromolecular complexes. Specifically, we identify RNP complexes that 1) dissociate, 2) form stable protein-only complexes, and 3) change composition in the absence of RNA suggesting specific roles for RNA in each of these cases. Because DIF-FRAC is independent of UV crosslinking, nucleotide incorporation, genetic manipulation or poly(A) RNA capture efficiency, it can therefore be used to investigate a wide variety of cell types, tissues and species. To demonstrate this versatility, we apply DIF-FRAC to mouse embryonic stem cells (mESCs), identifying 1,165 RNA-associated proteins, to show the method is highly adaptable and can be extended to discover RNP complexes in diverse samples.

Finally, we created a system-wide resource of 1,428 RNP complexes, many of which are previously unreported as having an RNA component, representing 20% of known human protein complexes. We provide our resource, rna.MAP, to the community as a fully searchable web database at rna.proteincomplexes.org.

## Results and Discussion

### <u>Dif</u>ferential <u>frac</u>tionation (DIF-FRAC) identifies RNP complexes

The DIF-FRAC strategy builds upon our previous strategy of Co-Fractionation Mass-spectroscopy (CF-MS) for identifying protein complexes in cellular lysate (Havugimana et al., 2012; Wan et al., 2015). CF-MS chromatographically separates protein complexes into fractions and uses a mass spectrometry pipeline to identify resident proteins in each fraction. The chromatographic elution profile of each protein is correlated to elution profiles from other proteins and similar profiles suggest physical interactions. Likewise, the DIF-FRAC strategy detects RNP complexes by identifying changes in the CF-MS elution profile of a protein complex’s subunits upon degradation of RNA (Figure 1).

We applied DIF-FRAC to human HEK 293T cell lysate using size-exclusion chromatography (SEC) to separate the cellular proteins in a control and an RNase A-treated sample into 50 fractions (Figure 1A). Upon RNase A treatment, we observed a loss in the bulk chromatography absorbance signal in the high-molecular-weight regions and an increase in absorbance in lower molecular weight regions, consistent with higher-molecular-weight species (>1000 kDa) becoming lower-molecular-weight species in the absence of RNA (Figure 1B). The distribution of cellular RNA in these fractions measured using RNA-seq confirmed that we are accessing a diverse RNA landscape of mRNAs, small RNAs, and lncRNAs (Figure S1). As a negative control, we applied the same DIF-FRAC strategy to human erythrocytes, which we reasoned should have fewer RNPs since they have substantially lower amounts of RNA due to the loss of their nucleus and ribosomes upon maturation (Keerthivasan et al., 2011). Accordingly, the absorbance chromatography signal of erythrocyte lysate showed only a negligible difference in a DIF-FRAC experiment (Figure 1C). Together, these data establish that DIF-FRAC is capable of identifying bulk changes to the RNA-bound proteome.

We next used mass spectrometry to identify and quantify the resident proteins in each fraction for both the control and RNase A treated chromatographic separations, resulting in 8,946 protein identifications. Using these abundance measurements, we compared elution profiles (i.e. abundance change across chromatographically-separated molecular weights) between the control and RNase A treated experiments for each protein. A shift in a protein’s elution profile between experiments is indicative of a protein-RNA interaction. For example, the known RNA helicase DDX21 shows a substantial shift in its elution profile upon RNase A treatment (Figure 1D), consistent with DDX21’s known association with RNA (Calo et al., 2015). Alternatively, proteins such as the glucose synthesis enzyme, PGM1, show no shift, consistent with it not binding RNA (Figure 1E).

We can further examine these elution profile differences in the context of physically-associated proteins to identify RNP complexes. For example, subunits of the spliceosome, a known RNP complex, show elution profiles that coelute in the control but shift markedly upon RNA degradation (Figure 1F). In contrast, the elution profiles of subunits of the non-RNA-associated hexameric MCM complex (M_r_ ∼550 kDa) (Figure 1G), as well as the 8-subunit COP9 signalosome (M_r_ ∼500 kDa) (Figure S2A), are unchanged by RNase A treatment, consistent with the complexes not interacting with RNA. Thus, DIF-FRAC produces a robust signal that can be used to differentiate between non-RNA-associated complexes and RNP complexes.

### Systematic identification of RNP complexes

In order to systematically identify RNP complexes in a DIF-FRAC experiment, we first developed a computational framework to identify statistically significant changes in individual proteins’ elution behavior to identify RNA-associated proteins. We observed a variety of changes in elution behavior of known RNA-associated proteins upon RNase A treatment including decrease in molecular weight (e.g. NCL), increase in molecular weight (e.g. SUGP1), decrease in observed abundance (e.g. RPL18A), and increase in observed abundance (e.g. MACF1) (Figure 1H). To capture this range of behaviors in a simple metric, we developed the “DIF-FRAC score,” which evaluates the degree to which two chromatographic separations differ (Figure 1I). Briefly, the DIF-FRAC score is a normalized Manhattan-distance between a protein’s control and RNase A treated elution profiles (see Methods). To identify significant changes, we calculated P-values by comparing each protein’s DIF-FRAC score to an abundance-controlled background distribution of DIF-FRAC scores from non-RNA-associated proteins (Figure S3A, see Methods for full description). We evaluated the score’s performance on a curated set of known RNA-associated proteins and see strong correspondence between precision and high-ranking proteins (Figure 1J, Figure S3B). DIF-FRAC identifies 1012 proteins with significant elution profile differences in HEK 293T cells with a false discovery rate (FDR) cutoff of 5% (Table S1). To validate our metric, our set of statistically significant hits was compared to RNA-associated proteins identified from 11 other studies using alternative methods including RNA interactome capture (RIC) (Baltz et al., 2012; Castello et al., 2012), organic phase separation (Queiroz et al., 2019; Trendel et al., 2019), and others (Hentze et al., 2018). These results indicate that the DIF-FRAC score is highly accurate for identifying individual RNA-associated proteins (Figure S4A-K).

**Table 1:**
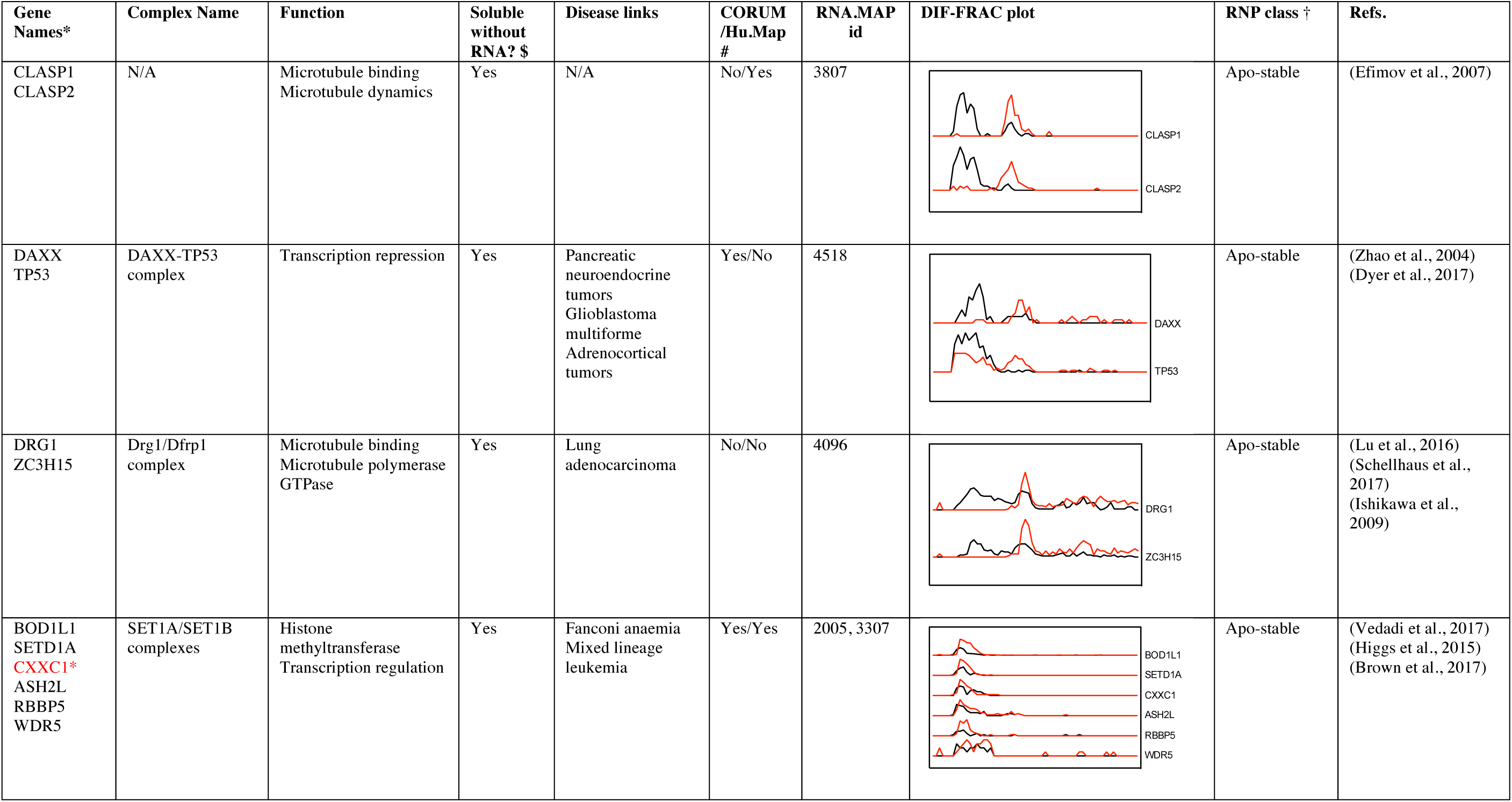
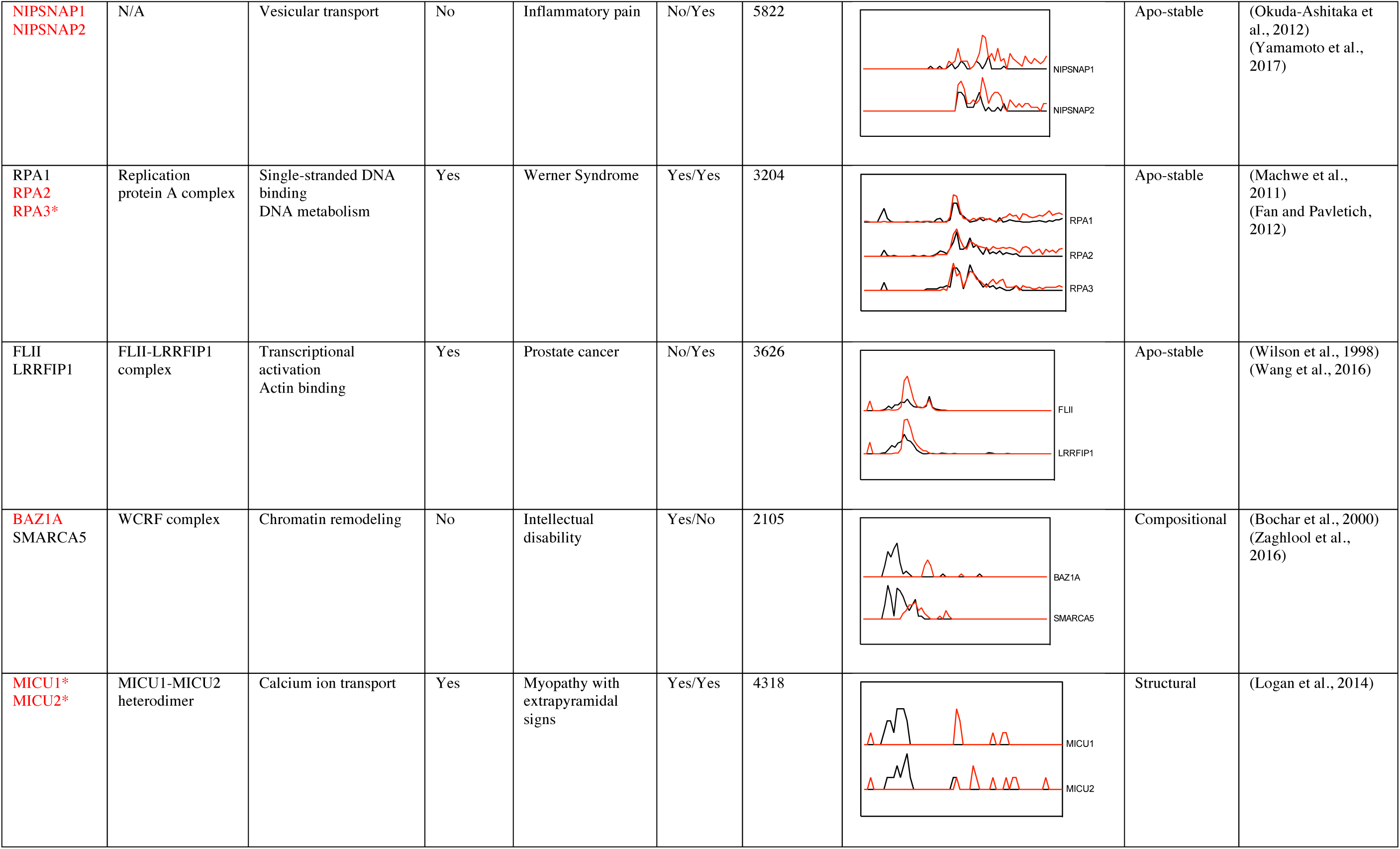
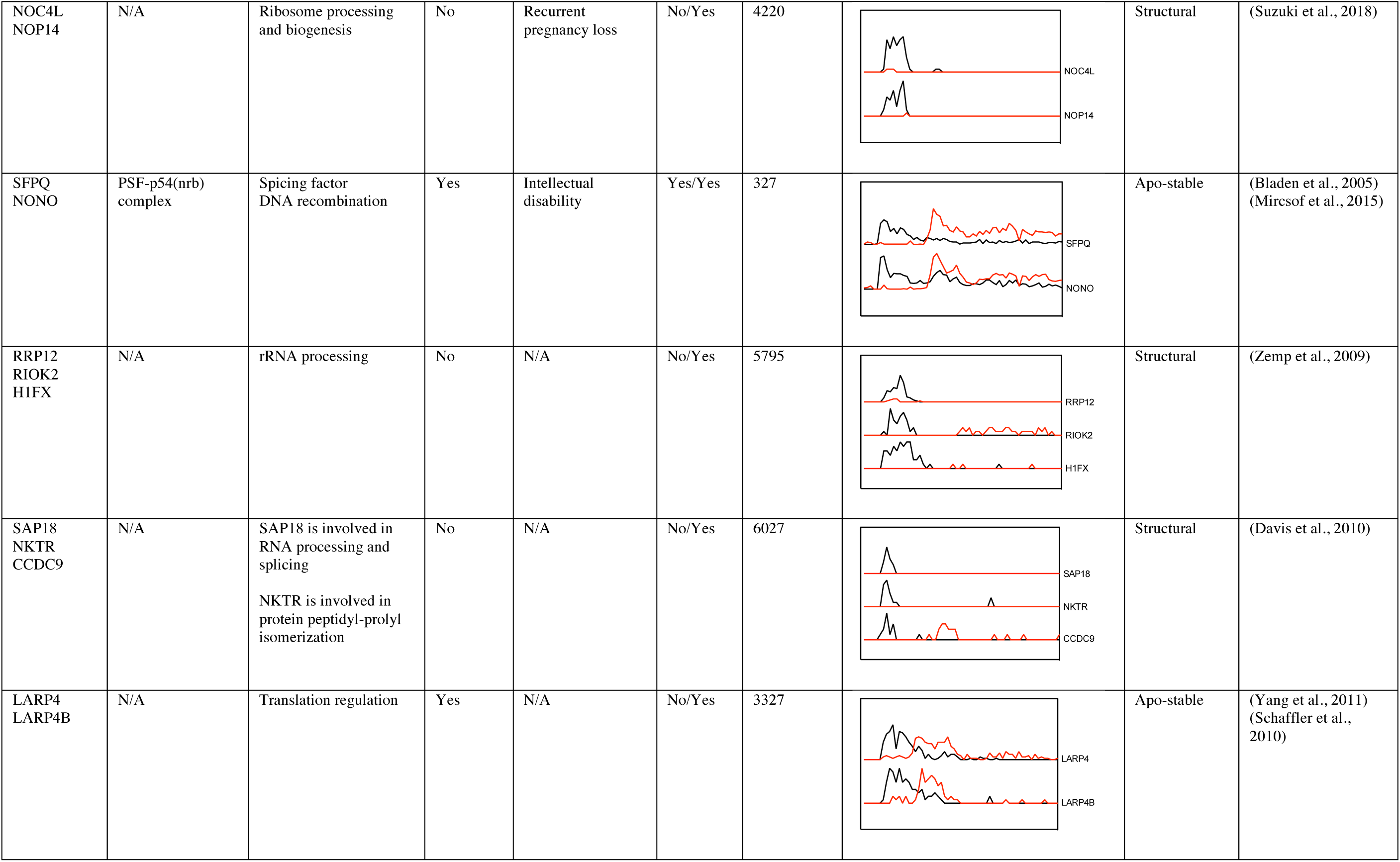
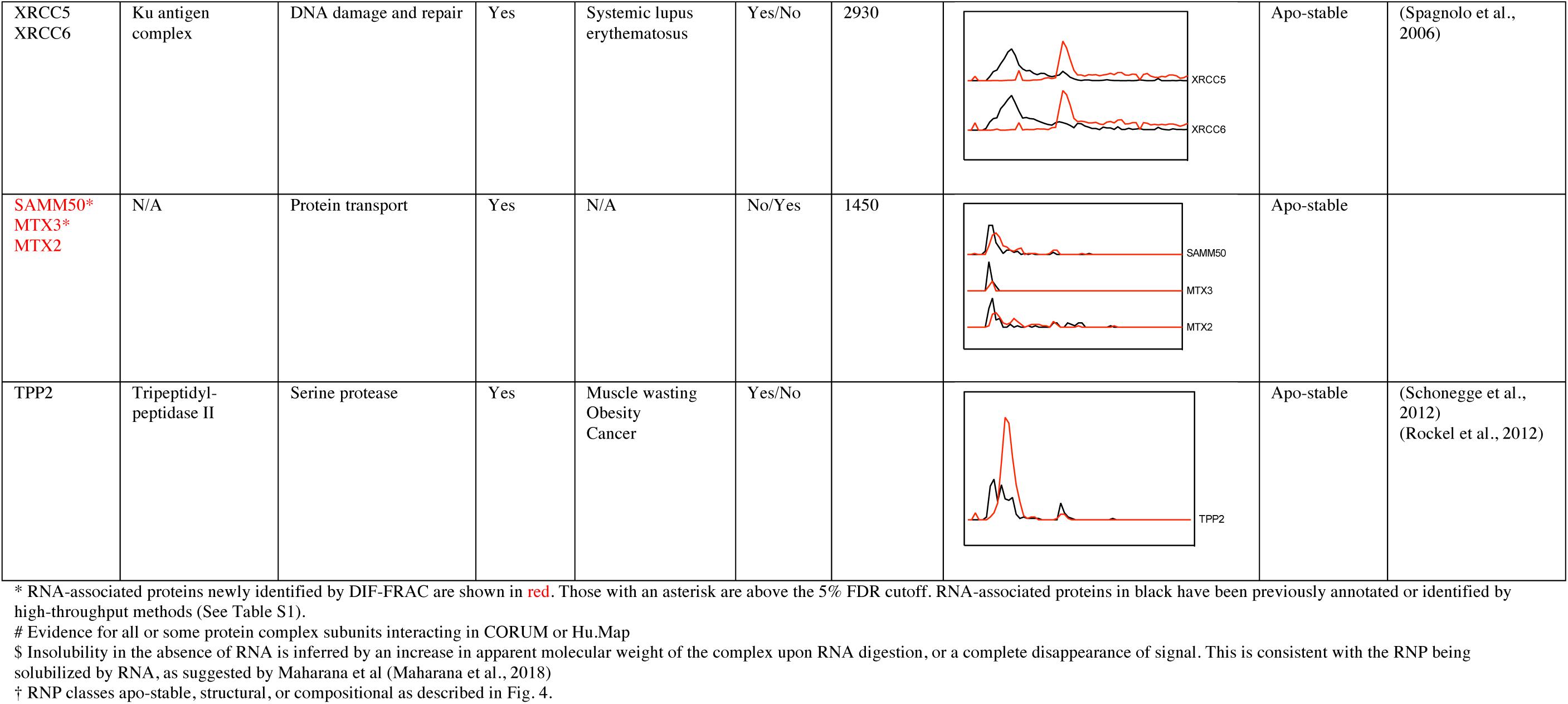
Stable RNPs identified by DIF-FRAC

To expand on previous systematic studies of RNA-associated proteins, we exploited the unique features of DIF-FRAC to identify which RNA-associated proteins are assembled into higher order RNP complexes. Specifically, we searched for protein complexes whose subunits co-elute in the control experiment in addition to being sensitive to RNase A treatment (e.g. see Figure 1F). We detected 115 RNP complexes that fit these criteria, which we term ‘RNP Select’ (Figure 2 and Table S2)). The RNP Select set consists of 464 unique proteins, and importantly, it recapitulates many known RNP complexes. The set includes canonical RNPs such as the 40S ribosome (Figure S5) and the spliceosomal tri-snRNP complex, a major component of the catalytically active spliceosome that contains an intricate network of snRNA binding interactions (Agafonov et al., 2016) (Figure 2B). The set also includes RNPs with more specific functions such as the IGF2BP1 complex, which stabilizes c-myc RNA and prevents translation-coupled decay (Weidensdorfer et al., 2009) (Figure 2B). Because these data demonstrated the veracity of the DIF-FRAC strategy, we next searched our dataset for additional insights into RNP biology.

**Figure 2:**
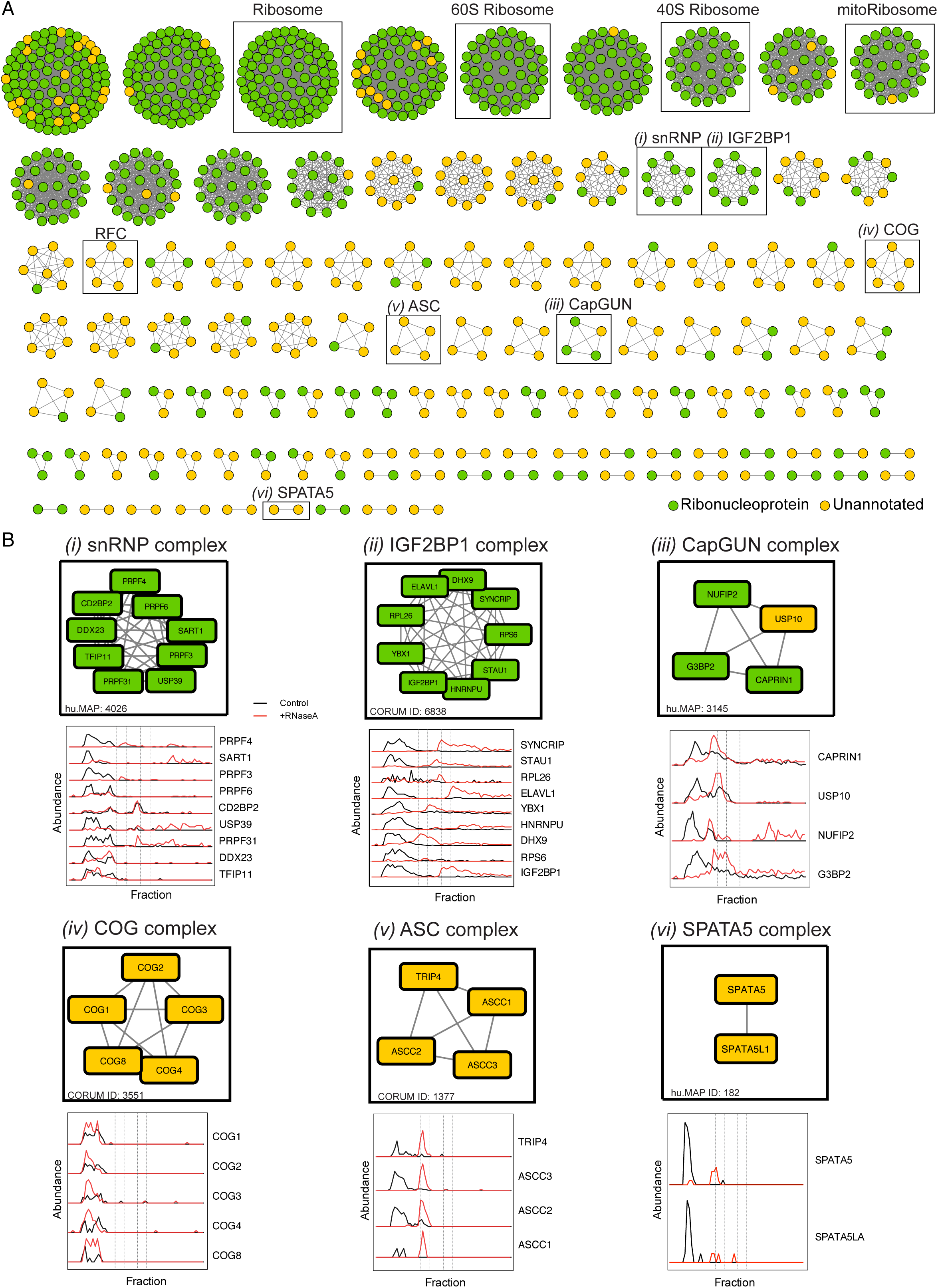
DIF-FRAC reveals a map of stable RNP complexes. (A) 115 RNP complexes identified by the DIF-FRAC method termed ‘RNP Select’. Green nodes represent proteins annotated as ‘ribonucleoprotein complex’ and yellow nodes are unannotated proteins. RNP Select complexes are defined as complexes whose protein subunits co-elute in the control DIF-FRAC sample (> 0.75 average correlation coefficient) and > 50% of subunits have a DIF-FRAC P-value > 0.5. DIF-FRAC identified many known RNP complexes such as the ribosome, mitochondrial ribosome and snRNP as well as novel RNP complexes such as RFC, COG, ASC and SPATA5. (B) Individual RNP complexes with elution profiles including *(i)* snRNP, *(ii)* IGF2BP1, *(iii)* CapGUN, *(iv)* COG, *(v)* ASC, and *(vi)* SPATA5. Abundance represents count of unique peptide spectral matches for each protein.

First, the RNP Select set provided new details about known RNPs. For example, stress granules are large membrane-less organelles that sequester mRNAs and prevent translation (Lin et al., 2015; Mittag and Parker, 2018; Spector and Lamond, 2011), and contain RNA-associated proteins including CAPRIN1, G3BP2, USP10 and NUFIP2 each localize to stress granules (Bardoni et al., 2003; Matsuki et al., 2013; Solomon et al., 2007). Interestingly, our previous map of human protein complexes (Drew et al., 2017) suggests that the known complex of G3BP, CAPRIN, and USP10 (Kedersha et al., 2016) also physically interacts with NUFIP2, leading us to suggest the name CapGUN (i.e. <u>CAP</u>RIN1, <u>G</u>3BP2, <u>U</u>SP10, <u>N</u>UFIP2). Importantly, DIF-FRAC revealed that CapGUN subunits co-elute and associate with RNA (Figure 2B).

More importantly, RNP Select also contains several complexes not previously known to associate with RNA. For example, the spinal muscular atrophy associated activating signal cointegrator (ASC) complex (Knierim et al., 2016) (Figure 2B) is a transcriptional coactivator of nuclear receptors and has a role in transactivation of serum response factor (SRF), activating protein 1 (AP-1), and nuclear factor kappaB (NF-kappaB) (Jung et al., 2002). Upon RNase A treatment, we observed a substantial shift in elution from a high molecular weight to a lower molecular weight for all subunits of the ASC complex strongly suggesting that the complex associates with RNA (Figure 2). Interestingly, one ASC component, ASCC1, has a predicted RNA binding motif near its C terminus and has been shown to localize to nuclear speckles (Soll et al., 2018), which like stress granules are membraneless organelles enriched for RNPs. Our results, in coherence with previous studies point to a role for the ASC complex associating with RNA in RNP granules. Other notable examples of previously uncharacterized RNP complexes include the conserved oligomeric Golgi (COG) complex which is involved in intra-Golgi trafficking, and the SPATA5-SPATA5L1 complex, an uncharacterized complex linked to epilepsy, hearing loss, and mental retardation syndrome (Tanaka et al., 2015) (Figure 2B) among others (Table 1).

Finally, to ascertain the total number of annotated protein complexes that likely function with an RNA component, we evaluated DIF-FRAC evidence for RNA-associated proteins in addition to the 11 other studies (described above) and identify 1,428 complexes that contain a majority of RNA-associated proteins (see Methods). This analysis suggests that greater than 20% of known protein complexes associate with RNA (Table 1 and Table S2). We provide the complete set of RNP complexes as a fully searchable web database, rna.MAP, at http://rna.proteincomplexes.org. This represents a detailed resource of human RNP complexes, providing myriad testable hypotheses to guide further explorations of RNP biology.

### Classification of RNP Complexes

RNA performs a variety of roles in macromolecular complexes. For example, it can bind as a substrate, function as an integral structural component, or act as a regulator of a complex’s composition. Mirroring these roles, DIF-FRAC data reveals that upon RNA degradation, the proteins in RNP complexes can remain in an intact complex (Figure 3A), become destabilized (Figure 3B), or adopt different higher order configurations (Figure 3C). We therefore categorize RNP complexes into three groups.

**Figure 3:**
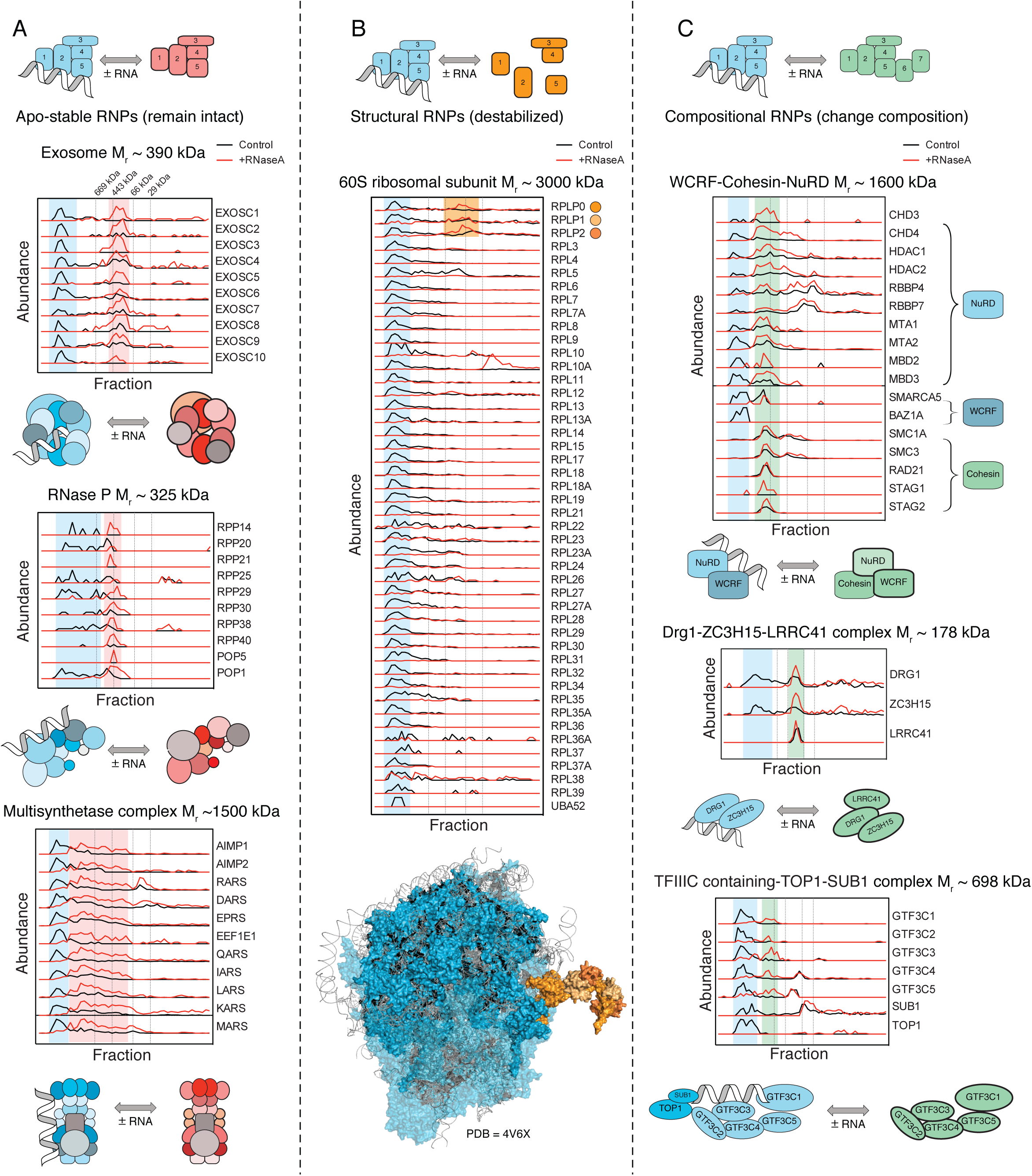
DIF-FRAC identifies 3 classes of RNP complexes. (A) ‘Apo-stable’ RNP complexes: elution profiles of the exosome (top, CORUM 7443), RNase P (middle, CORUM 123), and the multi-synthetase complex (bottom, CORUM 3040) show each complex is a stable complex that binds RNA and the complex remains intact in the absence of RNA. Blue shading represents RNA bound complex. Blue shading represents RNA bound form and red shading represents RNA unbound complex. (B) ‘Structural’ RNP complexes: elution profiles of the 60S ribosomal subunit (CORUM 308) show the complex destabilizes upon RNA degradation and subunits no longer co-elute upon RNase A treatment. DIF-FRAC elution data show the ribosomal subunits RPLP0, RPLP1 and RPLP2 (orange) remain as a subcomplex upon RNA degradation, consistent with their position in the solved ribosome structure whose interactions are not mediated by RNA (bottom, PDB 4V6X, protein in blue, RNA in red, ribosomal stalk in orange). (C) ‘Compositional’ RNP complexes: (Top) Elution profiles of WCRF-Cohesin-NuRD (CORUM 282) and NuRD-WCRF suggest that RNA association promotes different forms of the complex. (Middle) Elution profiles of Drg1-ZC3H15-LRRC41 complex (hu.MAP 2767) which forms only in the absence of RNA. (Bottom) Elution profiles of the TFIIIC containing-TOP1-SUB1 complex (CORUM 1106) loses two subunits, TOP1 and SUB1, upon RNA degradation. Green shading represents RNA unbound complex. In (A)-(C) vertical dashed lines correspond molecular weight standards described in Figure 1.

The first category, which we term “apo-stable,” defines protein complexes that remain stable after RNase A treatment. These include the exosome, RNase P, and the multi-synthetase complex (Figure 3A). Elution profiles of apo-stable complexes show that in the absence of RNA, subunits still co-elute, but do so as a lower-molecular weight complex. Available atomic structures of the exosome with and without RNA support the concept that RNA is peripheral to the stability of the complex (Gerlach et al., 2018; Weick et al., 2018).

The second category, which we designate as “structural”, refers to complexes for which RNA is essential for the RNP complex structure and/or subunit solubility. These include, for example, the 60S and 40S ribosomal subcomplexes (Figure 3B and Figure S5). Upon degradation of RNA, the observed abundance of ribosomal protein subunits markedly decreases; suggesting the ribosome breaks apart and subunits become insoluble. This result is consistent with solved structures of the ribosome (Anger et al., 2013), demonstrating the centrality of rRNAs to the overall complex architecture (Figure 3B). Interesting exceptions to this behavior are the DIF-FRAC elution profiles for RPLP0, RPLP1 and RPLP2. These proteins co-elute in the RNase A treated sample, suggesting RNA does not mediate their interaction. Strikingly, however, this observation is consistent with the atomic structure of the human ribosome, which suggests that interactions between RPLP0, RPLP1 and RPLP2 are entirely protein-mediated (Figure 3B). This example demonstrates how DIF-FRAC data can not only identify RNA-protein mediated interactions, but can also provide structural information about RNP subcomplexes.

The third category, ‘compositional’ complexes, refers to those in which RNA promotes different stable combinations of protein-complex subunits, perhaps in a regulatory role (Figure 3C). For example, the WCRF (Williams syndrome transcription factor-related chromatin remodeling factor) complex, NuRD (Nucleosome Remodeling Deacetylase) complex and Cohesin complex are reported to assemble into a chromatin-remodeling supercomplex (CORUM ID: 282). We observed the WCRF and NuRD complexes co-eluting in the control experiment, forming a 12-subunit complex that shifts its elution upon RNA degradation. Interestingly, we also observed the supercomplex (WCRF, NuRD and Cohesin) eluting as a ∼17-subunit complex in the RNA degradation condition. This composition change provides an explanation for why several NuRD-containing complexes are observed experimentally (Hakimi et al., 2002; Xue et al., 1998); our data suggest that these may represent both RNP complexes and non-RNA-associated complexes.

We also identified an uncharacterized compositional RNP complex containing the cell growth regulators DRG1 and ZC3H15 (DRFP1) (Ishikawa et al., 2005) that are implicated in lung cancer (Lu et al., 2016). ZC3H15 stabilizes DRG1 and prevents degradation possibly by preventing poly-ubiquitination (Ishikawa et al., 2005). Our result suggests that RNA is also involved in ZC3H15’s role in stabilizing DRG1, as we observed a shift to a non-RNA-associated complex containing DRG1-ZC3H15 and LRRC41 in the absence of RNA (Figure 3C). LRRC41 is a probable substrate recognition component of E3 ubiquitin ligase complex (Kamura et al., 2004).

A further example of a compositional RNP complex is the transcription factor (TF)IIIC-TOP1-SUB1 complex, which is involved in RNA polymerase III pre-initiation complex (PIC) assembly (Male et al., 2015). DIF-FRAC shows this 7-subunit complex changes composition to the five-subunit TFIIIC upon RNA degradation (Figure 3C), offering further insights into the mechanism of TFIIIC-dependent PIC formation.

Finally, we identified the chromatin remodeling BRG/hBRM associated factors (BAF; the mammalian SWI/SNF complex; SWI/SNF-A) and polybromo-associated BAF (PBAF; SWI/SNF-B) complexes as compositional RNP complexes, which is significant because these are some of the most frequently mutated protein complexes in cancer (Hodges et al., 2016; Tang et al., 2017) (Figure S6). BAF and PBAF complexes share a set of common core subunits, but also each have signature subunits that are related to their respective functions. Elution profiles show these core subunits co-elute with PBAF-only subunits in the control, but co-elute with BAF-only subunits upon RNA degradation (Figure S6). These data suggest BAF exists as a non-RNA-associated complex while PBAF functions as an RNP complex, consistent with its known role in transcription and supporting a previously described RNA-binding model where lncRNAs interact with SWI/SNF complexes in cancer (Tang et al., 2017). Together, these examples demonstrate the power of DIF-FRAC to describe the various physical relationships between RNA and macromolecular protein complexes.

### Characterization of individual RNA-associated proteins

Although our efforts focused primarily on higher order RNP complexes, it is important to note that DIF-FRAC is also a powerful complement to existing methods for characterizing individual RNA-associated proteins. Indeed, DIF-FRAC identified 196 human RNA-associated proteins not previously identified in the many previous studies discussed in the Introduction, above (Table S3, Figure S4L). These DIF-FRAC identified RNA-associated proteins were strongly enriched in RNA binding domains annotated by Interpro (Finn et al., 2017) (Figure S4M). As we described above, inspection of elution profiles for the individual proteins revealed at least four distinct DIF-FRAC signals (Figure 1H, Figure 4). These manifest as elution-profile shifts with RNase A treatment that show: (1) an apparent decrease in molecular weight of the RNA-associated protein consistent with the degradation of an RNA component (Figure 4A); (2) a decrease in observed abundance, suggesting the RNA-associated protein becomes insoluble or is degraded (Figure 4B); (3) an apparent increase in molecular weight, suggesting the RNA-associated protein forms a higher-order species or aggregate (Figure 4C); or (4) an increase in observed abundance, indicative of the RNA-associated protein becoming more soluble (Figure 4D).

**Figure 4:**
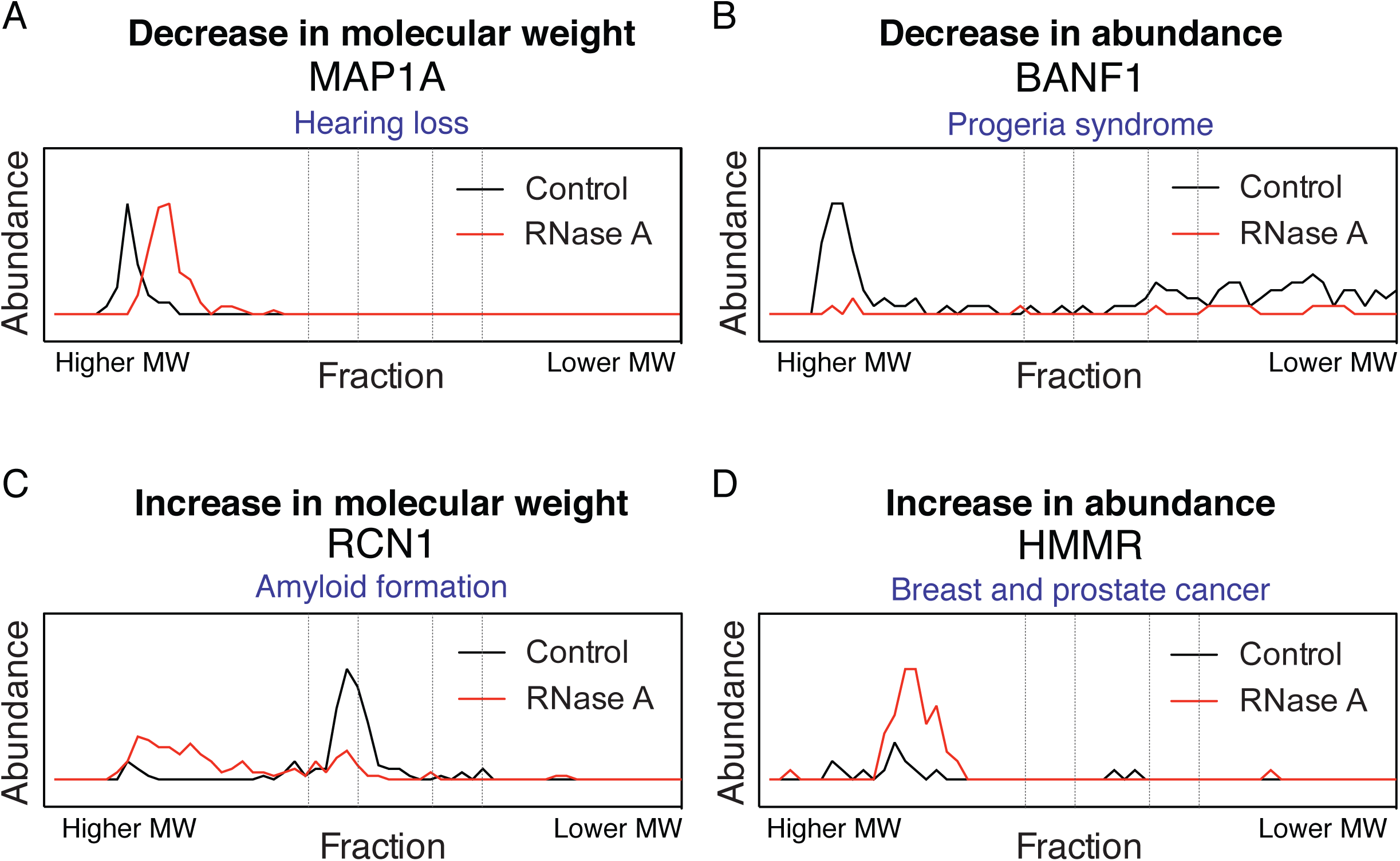
DIF-FRAC identifies four distinct signals for RNA-associated proteins. Examples of elution profiles for disease related proteins that (A) decrease in size, MAP1A, (B) decrease in observed abundance (less soluble), BANF1, (C) increase in size, RCN1, and (D) increase in observed abundance (more soluble), HMMR, upon RNA degradation.

Analysis of all identified RNA-associated proteins shows 796 (79%) decrease in molecular weight, while 216 (21%) RNA-associated proteins increase in size (Figure S7). Aside from RNA acting as an interaction partner to RNA-associated proteins, RNA has been shown to regulate the oligomerization state of proteins both positively (Bleichert and Baserga, 2010; Huthoff et al., 2009; Xie et al., 2018) and negatively (Yoshida et al., 2004). Our data suggests that while the majority of RNA-associated proteins form higher-order assemblies with RNA, the oligomerization of 21% is potentially inhibited by RNA. Alternatively, RNA has also been shown to alter the solubility state of proteins (Maharana et al., 2018). We observe an increase in observed abundance for 535 (53%) proteins upon RNase A treatment, a decrease in abundance for 470 (47%) proteins, and no change in observed abundance for only 7 proteins. This suggests RNA impacts the solubility for most RNA-associated proteins and may function to tune protein availability in the cell.

Looking specifically at individual proteins provided insights that could impact our understanding of human disease. For example, we found that BANF1, a chromatin organizer, appears insoluble under our experimental conditions without RNA (Figure 4B). Interestingly, the BANF1 mutation Ala12-Thr12 causes Hutchinson-Gilford progeria syndrome, a severe and debilitating aging disease, by a reduction in protein levels (Puente et al., 2011). Our data suggest the hypothesis that this reduction is caused by disruption of the RNA-BANF1 interaction, leading to insolubility and degradation. Furthermore, RNA has also been shown to solubilize proteins linked to pathological aggregates (Maharana et al., 2018). Our data identifies a number of CREC family members (CALU, RCN1, RCN2, and SDF4; Figure 4C and Table S1) as RNA-associated proteins that increase in molecular weight upon RNA degradation. The CREC family is a group of multiple EF-hand, low-affinity calcium-binding proteins with links to amyloidosis (Vorum et al., 2000). DIF-FRAC demonstrates a dependence of RNA on the oligomerization state of CALU, which could play a role in the formation of amyloid deposits similar to that observed for prion-like RNA-associated proteins (Maharana et al., 2018). Based on these examples and the many disease links to DIF-FRAC identified RNP complexes (Table 1), we anticipate our data will generate testable RNA-related hypotheses about disease-related states.

### Directed validation of Replication factor C (RFC) as an RNP Complex

An important aspect of DIF-FRAC is that while it provides a systematic survey, the experimental basis for each data point can be directly assessed in the elution profiles. Nonetheless, the ultimate demonstration of the utility of any large-scale dataset is its ability to make predictions that can be validated by orthogonal experiments. Among the most surprising findings in our data was that the extensively-characterized replication factor C (RFC) complex (Yao and O’Donnell, 2012) exists as a stable RNP complex (Figure 2A). During replication and DNA damage repair, the RFC complex is responsible for loading PCNA, a DNA polymerase processing factor, onto DNA. Strikingly, DIF-FRAC identified two previously observed variants of the RFC complex, RFC1-5 and RFC2-5 (Figure 5A), and more importantly demonstrated that RFC1-5 appears to be the dominant variant and is also the RNA-associated form (Figure 5C). Consistent with the RFC complex interacting with RNA, the homologous clamp loader in *E.coli*, γ complex, is known to load the DNA clamp onto RNA-primed template DNA (Yao and O’Donnell, 2012) and eukaryotic RFC has also been shown to be capable of loading PCNA onto synthetic RNA-primed DNA (Yuzhakov et al., 1999). In light of this finding, we tested whether purified RFC complex from *S. cerevisiae* could directly bind different species of nucleic acids. We observed that RFC not only binds dsDNA and DNA-RNA hybrids, but also binds dsRNA with surprisingly tight binding constants in the nanomolar range (Figure 5D and Figure S8). These data show RFC binds dsRNA and point to an uncharacterized role for RNA in the function of RFC. The result also further validates the use of DIF-FRAC to identify uncharacterized RNP complexes.

**Figure 5:**
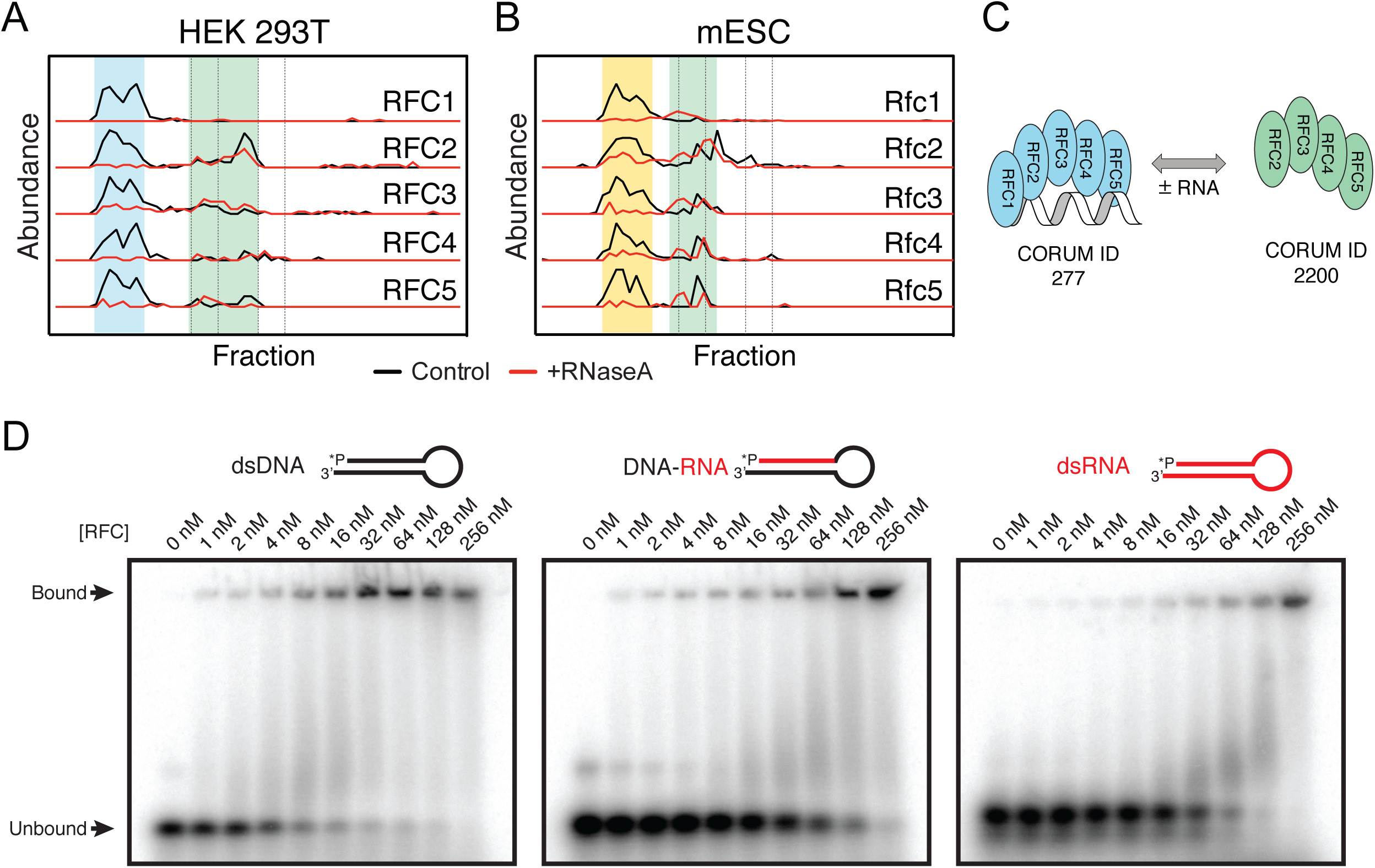
Replication factor C is an RNP Complex. Elution profiles in both human (A) and mouse (B) demonstrates RFC1-5 forms an RNP complex (blue/yellow highlight). A smaller subcomplex of RFC2-5 (green highlight) becomes the dominant form upon RNA degradation. (C) A cartoon to show the RNA-dependence of annotated complexes RFC1-5 (blue) and RFC2-5 (green) as determined by DIF-FRAC. RNA is shown in grey. (D) Electromorphic mobility shift assays (EMSA) of various concentrations of purified *S. cerevisiae* RFC mixed with 1 nM ^32^P-labeled oligonucleotides. Representative gels show RFC binds dsDNA, DNA/RNA hybrid and dsRNA substrates. RFC-nucleic acid complexes were separated on 10% native gels. Binding constants are in the nanomolar range (Figure S8).

### Evaluating RNPs in multiple proteomes

Finally, because DIF-FRAC does not rely on any specialized reagents, the strategy can be applied to any cell type that can be readily isolated. Because of the long-standing interest in the role of RNPs in embryonic development (e.g. for targeted localization of maternal RNAs (Escobar-Aguirre et al., 2017); processing of non-coding RNA to direct differentiation and stem cell potency (Dinger et al., 2008; Guttman et al., 2011; Yan et al., 2013)), we applied DIF-FRAC to mouse embryonic stem cells (mESCs). We identified 1,165 significant RNA-associated proteins in mESCs (Figure 6A, Table S1), including 466 that are novel, representing a 35% increase in the number of annotated mouse RNA-associated proteins (Figure 6B). This mESC dataset provides three advances:

**Figure 6:**
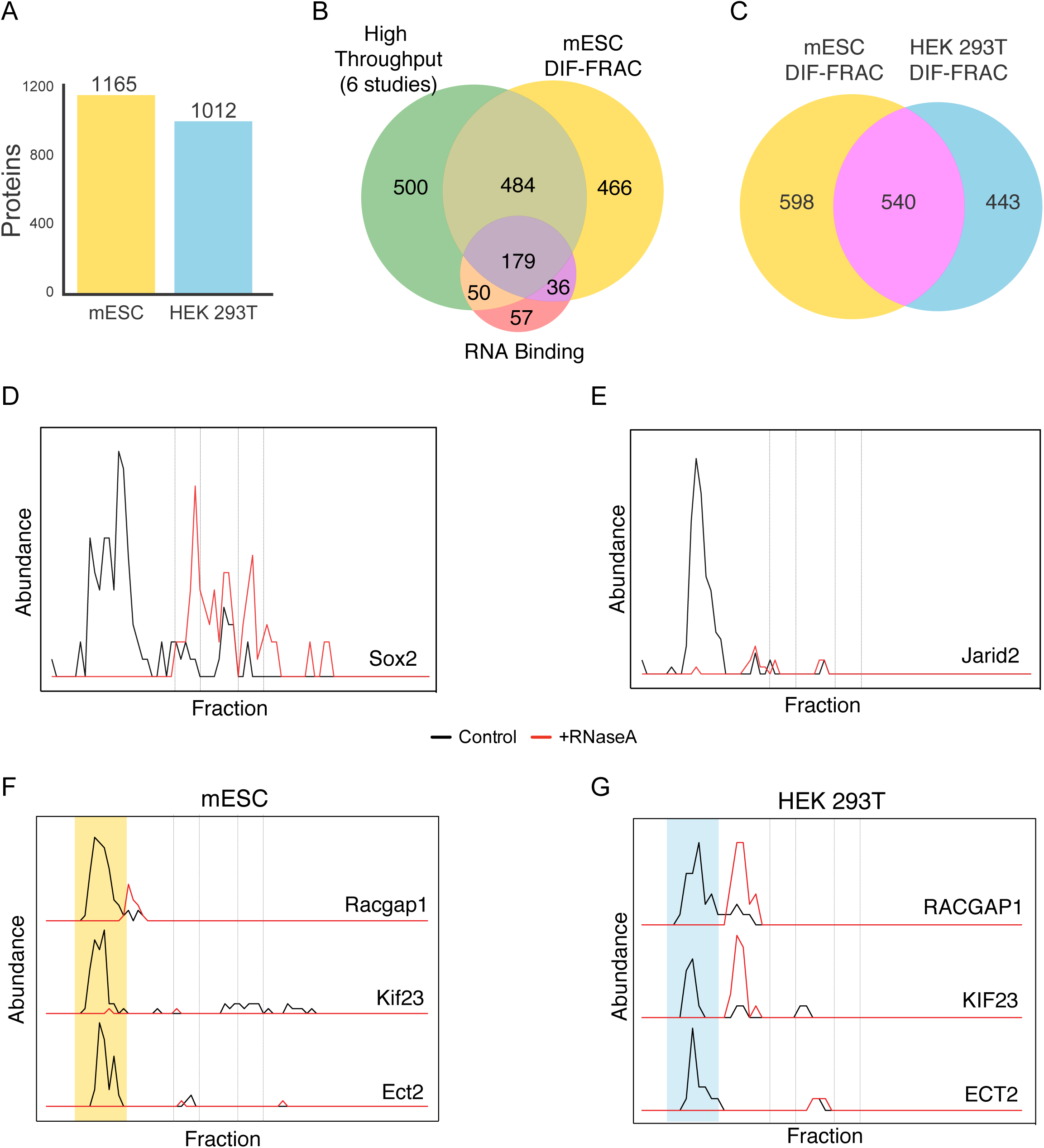
DIF-FRAC identifies RNP complexes across cell types and species. (A) DIF-FRAC identifies 1,165 RNA-associated proteins in mESC (mouse embryonic stem cells), and 1,012 RNA-associated proteins in HEK 293T cells. (B) Venn diagram of considerable overlap between previously published large-scale RNA-protein interaction studies, literature annotated RNA-associated proteins, and DIF-FRAC identified RNA-associated proteins in mESC. (C) RNA-associated human-mouse orthologues are identified reproducibly in DIF-FRAC experiments. Elution profiles for known pluripotency factors, Sox2 (D) and Jarid2 (E) show association with RNA in mESC cells. Elution profiles of the centralspindlin complex for (F) mESC and (G) HEK 293T demonstrate centralspindlin is an RNP complex in both species. Yellow and blue shading represent RNA bound complex in mESC and HEK 293T respectively.

First, the data can provide additional evidence to support assignment of novel RNPs. For example, many of the RNA-associated proteins identified in mESCs reflected equivalent RNA-associated proteins in human cells (Figure 6C), including the RFC complex, which specifically behaves as a compositional RNP complex in both species (Figure 5B).

Second, this approach allowed the identification of cell-type specific RNA-associated proteins. Indeed, we identified several mESC-specific RNA-associated proteins and these included several that have been previously implicated in stem cell function. For example, we identified the known pluripotency factor Sox2 (Figure 6D) and the Polycomb Repressor complex 2 subunit Jarid2 (Figure 6E), as RNA-associated proteins, consistent with previous reports (Cifuentes-Rojas et al., 2014; Fang et al., 2011; Kaneko et al., 2014).

Finally, the additional dataset allows us to cast a wider net in our search for novel RNPs. For example, among the RNA-associated proteins identified in mESCs were members of the centralspindlin complex, a heterotetramer consisting of Racgap1 and Kif23 and involved in cytokinesis (White and Glotzer, 2012; Yüce et al., 2005). Previously unknown to contain an RNA component, we identify Racgap1, Kif23 and the centralspindlin interaction partner Ect2 as significantly sensitive to RNase A treatment in mESCs (Figure 6F). In agreement with this mESC result, we observed a similar trend for this complex in human cells, showing conservation across species (Figure 6G). Our results suggest a physical interaction between the centralspindlin complex and RNA, thus informing a previous study that report Kif23 (ZEN-4 in *C.elegans*) as a positive regulator of RNP granule formation (Wood et al., 2016), as well as the localization of several RNA species to the midbody during cytokinesis (Clemson et al., 1996; Lécuyer et al., 2007; Zheng et al., 2010).

Together, these data demonstrate that the adaptability of DIF-FRAC to diverse systems will allow identification of conserved RNA-associated proteins and RNP complexes in diverse tissues and disease states across all domains of life.

### Conclusion

Here, we report the design, development, and application of a robust fractionation-based strategy to determine RNP complexes on a proteome-wide scale. We successfully used DIF-FRAC to identify 115 stable RNP complexes throughout the human interactome, and applied it to multiple tissue types and species. Combining this with previous data, we generate a resource of the RNA-bound human proteome and demonstrate that upwards of 20% of protein complexes contain an RNA component, highlighting the prevalence of RNP complexes in the cellular milieu. Together our results provide a valuable tool for researchers to investigate the role of RNPs in protein function and disease.

The DIF-FRAC methodology offers important advances over previous techniques that examine RNA-protein interactions. Specifically, interactions are probed proteome-wide in a native, whole lysate sample using a strategy that is not reliant on labeling or cross-linking efficiency. We show DIF-FRAC can be applied effectively to multiple cell types and organisms, and has the potential to provide information on protein-RNA interactions in disease states. Furthermore, DIF-FRAC is a broadly applicable framework that can be extended to examine other large-scale proteomic changes in a system of interest.

We also introduce three classifications of RNP complexes (apo-stable, structural, and compositional) that provide a useful framework to organize the roles of RNAs in macromolecular complexes. Additionally, DIF-FRAC provides information on the biochemical characteristics (i.e. molecular weight, solubility) of RNP complexes in the presence and absence of RNA that offer clues to disease pathophysiology. We anticipate this technique to be a powerful tool to uncover the molecular mechanisms of RNA related diseases. Overall, the DIF-FRAC method described and demonstrated here charters new territories in the cellular landscape of RNA-protein interactions. We have utilized DIF-FRAC to provide the first system-wide resource of human RNPs, providing a broadly applicable tool for studying cellular interactions and responses in multiple cell types and states.

## Acknowledgements

We would like to thank Matt Davis for helpful comments, Claire D. McWhite for plotting software and the Texas Advanced Computing Center for high-performance computing. This work was supported by grants from the NIH (K99 HD092613 to K.D., R35 GM122480, DP1 GM106408, R21 GM119021, R01 DK110520, and R01 HD085901 to E.M.M., R01 GM112722 to J.K. and R01 GM120554 to I.J.F.); NSF (IOS-1237975 to E.M.M., and 1453358 to I.J.F.); Army Research Office (W911NF-12-1-0390 to E.M.M.); and The Welch Foundation (F1515 to EMM and F1808 to I.J.F.). Data collection by the UT Austin Proteomics Facility was supported by CPRIT RP110782 to Maria Person.

## Materials and Methods

### Experimental Design

#### Human cell culture and extract preparation

HEK293T cells (ATCC CRL3216) cultured in DMEM (Gibco) supplemented with 10 % (v/v) FBS (Life Technologies) were continually split over 7 days to give four 10-cm dishes of adherent cells. For the control fractionation sample, two 10-cm dishes of cells were harvested at 80-100 % confluence without trypsin by washing in ice cold phosphate buffered saline (PBS) pH 7.2 (0.75 mL; Gibco) and placed on ice. Cells (approximately 0.1 g wet weight) were lysed on ice (5 min) by resuspension in Pierce IP Lysis Buffer (0.8 mL; 25 mM Tris-HCl pH 7.4, 150 mM NaCl, 1 mM EDTA, 1% NP-40 and 5% glycerol; Thermo Fisher) containing 1x protease inhibitor cocktail III (Calbiochem). The resulting lysate was clarified (17,000g, 10 min, 4°C) and left at room temperature (30 min). The sample was filtered (Ultrafree-MC filter unit (Millipore); 12,000g, 2 min, 4°C) to remove insoluble aggregates. RNase A treated samples were prepared on the same day in an identical manner, except RNase A (8 μL, 80 μg, Thermo Fisher, catalogue #EN0531) was added after lysate clarification and the sample left at room temperature (30 min) before filtration.

#### Mouse embryonic stem cell culture

Gelatin adapted mouse J1 ES cells (ATCC® SCRC-1010™) were cultured in Dulbecco’s Modified Eagle’s Medium (DMEM, Life Technologies) containing 18% fetal bovine serum (FBS, Gemini), 50 U/mL of penicillin/streptomycin with 2 mM L-glutamine (Life Technologies), 0.1 mM non-essential amino acid (Life Technologies), 1% nucleosides (Sigma-Aldrich), 0.1 mM β-mercaptoethanol (Sigma-Aldrich), and 1,000 U/mL recombinant leukemia inhibitory factor (LIF, Chemicon). ES cells were plated on 15-cm dishes coated with 0.1% gelatin and incubated at 37°C and 5% CO_2_. Cells were passaged every 2 days. Lysis and RNase A treatment were done as described in the HEK 293T protocol.

#### Erythrocyte cell preparation

Leukocyte-reduced red blood cells (RBCs) were obtained from an anonymous donor and purchased from Gulf Coast Regional Blood Center (Houston, Texas). The RBCs used in this experiment were kept at 4°C for 54 days before lysis to ensure reticulocytes mature into RBCs. Prior to cell lysis, RBCs were washed with ice cold PBS (pH 7.4, Gibco) for 3 times at 600 g for 15 min at 4°C. RBCs were then lysed in hypotonic solution (5 mM Tris-HCl, pH 7.4) containing protease and phosphatase inhibitors (cOmplete, EDTA-free Protease Inhibitor Cocktail, Roche and PhosSTOP, Roche) with a ratio of 1 volume packed RBC: 5 volumes hypotonic solution. Hemolysate (soluble fraction of RBC lysate) was collected by centrifuging white ghosts (membrane fraction of RBC lysate) at 21,000 g for 40 mins at 4°C. Hemolysate was collected and stored at −80°C until further use. On the day of experiment, hemolysate was thawed and treated with Hemoglobind (Biotech Support Group) in order to remove hemoglobin from hemolysate. A total of 4-5 mg of total proteins were split into control and RNase A treated samples. The RNase sample was treated with RNase A as described in the protocol of RNase A treatment of lysate from HEK293T cells. Both samples were filtered (Ultrafree-MC filter unit (Millipore); 12,000 g, 2 min, 4°C) to remove insoluble aggregates prior to fractionation.

#### Biochemical fractionation using native size-exclusion chromatography

All lysates were subject to size exclusion chromatography (SEC) using an Agilent 1100 HPLC system (Agilent Technologies, ON, Canada) with a multi-phase chromatography protocol as previously described (Havugimana et al., 2012). Soluble protein (1.25 mg, 250 μL) was applied to a BioSep-SEC-s4000 gel filtration column (Phenomenex) equilibrated in PBS, pH 7.2 (HEK 293T and mESC lysate) or pH 7.4 (erythrocytes) at a flow rate of 0.5 mL min^-1^. Fractions were collected every 0.375 mL. The elution volume of molecular weight standards (thyroglobulin (M_r_ = 669 kDa); apoferritin (M_r_ = 443 kDa); albumin (M_r_ = 66 kDa); and carbonic anhydrase (M_r_ = 29 kDa); Sigma) was additionally measured to calibrate the column (Figure 1B).

#### Mass spectrometry

Fractions were filter concentrated to 50 μL, denatured and reduced in 50 % 2,2,2-trifluoroethanol (TFE) and 5 mM tris(2-carboxyethyl)phosphine (TCEP) at 55 °C for 45 minutes, and alkylated in the dark with iodoacetamide (55 mM, 30 min, RT). Samples were diluted to 5 % TFE in 50 mM Tris-HCl, pH 8.0, 2 mM CaCl_2_, and digested with trypsin (1:50; proteomics grade; 5 h; 37 °C). Digestion was quenched (1 % formic acid), and the sample volume reduced to ∼100 μL by speed vacuum centrifugation. The sample was washed on a HyperSep C18 SpinTip (Thermo Fisher), eluted, reduced to near dryness by speed vacuum centrifugation, and resuspended in 5 % acetonitrile/ 0.1 % formic acid for analysis by LC-MS/MS.

Peptides were separated on a 75 μM x 25 cm Acclaim PepMap100 C-18 column (Thermo) using a 3-45 % acetonitrile gradient over 60 min and analyzed online by nanoelectrospray-ionization tandem mass spectrometry on an Orbitrap Fusion or Orbitrap Fusion Lumos Tribrid (Thermo Scientific). Data-dependent acquisition was activated, with parent ion (MS1) scans collected at high resolution (120,000). Ions with charge 1 were selected for collision-induced dissociation fragmentation spectrum acquisition (MS2) in the ion trap, using a Top Speed acquisition time of 3-s. Dynamic exclusion was activated, with a 60-s exclusion time for ions selected more than once. MS from HEK 293T cells was acquired in the UT Austin Proteomics Facility.

#### Construction and sequencing of RNA-seq libraries of DIF-FRAC samples

Fractions from a biological replicate SEC separation corresponding to higher molecular weight species (approximately >1.5 MDa; fractions 16-23 in Figure 1B) were analyzed by total RNA sequencing. Total RNA was isolated from each fraction (0.375 mL) by addition of Trizol (1.125 mL; Thermo Fisher) and the sample (1.4 mL) was transferred to a Phasemaker tube (Thermo Fisher). Total RNA was extracted following the protocol supplied by the manufacturer and further cleaned up using a RNeasy MinElute Cleanup Kit (Qiagen). RNA integrity number (RIN) was measured using an Agilent Bioanalyzer and samples were ribo-depleted using a using a RiboZero Gold (Human/Mouse/Rat) kit (Illumina) to remove rRNAs. RNA libraries were prepared for sequencing according to vendor protocols using NEBNext R Small RNA Library Prep Set for Illumina R (Multiplex Compatible), Cat #E7330L, according to the protocol described by Podnar et al. (Podnar et al., 2014). RNA was fragmented using elevated temperature in carefully controlled buffer conditions to yield average fragment sizes of 200 nucleotides. These fragments were directionally ligated to 5′ and 3′ adaptors so that sequence orientation is preserved throughout sequencing. Reverse transcription and PCR were performed to complete the DNA sequencing libraries, which were sequenced using an Illumina NextSeq 500 instrument (75-nt single reads) at the Genomic Sequencing and Analysis Facility at the University of Texas at Austin.

#### *S. cerevisiae* RFC purification

RFC was purified as previously described (Finkelstein et al., 2003; Kim et al., 2017). Briefly, full-length *S. cerevisiae* RFC was expressed in BL21(DE3) ArcticExpress (Agilent) *E. coli* co-transformed with pLant2b-RFC-AE (pIF117) and pET11-RFC-BCD (pIF116). RFC was subsequently purified by SP and Q (GE Healthcare) ion exchange chromatography. Protein concentration was determined by comparison to a BSA titration curve using Coomassie-stained SDS-PAGE.

#### Electrophoretic Mobility Shift Assay (EMSA)

Oligonucleotide constructs were based on an earlier description (Kobayashi et al., 2006). Each of the four-nucleic acid substrates were radiolabeled with [γ-^32^P]-ATP using T4 Polynucleotide Kinase (NEB). Free nucleotide was removed using G-25 MicroSpin columns (GE Healthcare). Oligonucleotides were subsequently heated to 75°C and slowly cooled to room temperature to allow proper annealing. 1 nM oligonucleotide and various concentrations of RFC (0 to 256 nM) were incubated for 15 minutes at room temperature in a buffer containing 25 mM Tris-HCl [pH 7.5], 50 mM NaCl, 2 mM MgCl_2_, 2 mM DTT, and 0.1 mg/mL BSA. Reactions were quenched with 6x loading dye (10 mM Tris-HCl [pH 7.6], 60% glycerol, 60 mM EDTA, 0.15% [w/v] Orange G) and subsequently separated by native acrylamide gel electrophoresis. Gels were dried on Zeta-Probe Membrane (Bio-Rad) at 80°C for two hours. Bands were visualized by a Typhoon FLA 7000 phosphorimager (GE Healthcare). Binding was quantified using FIJI (Schindelin et al., 2012). Subsequent data were fit to a hyperbolic equation to determine the k_D_ for oligonucleotide binding.

#### Oligonucleotides used

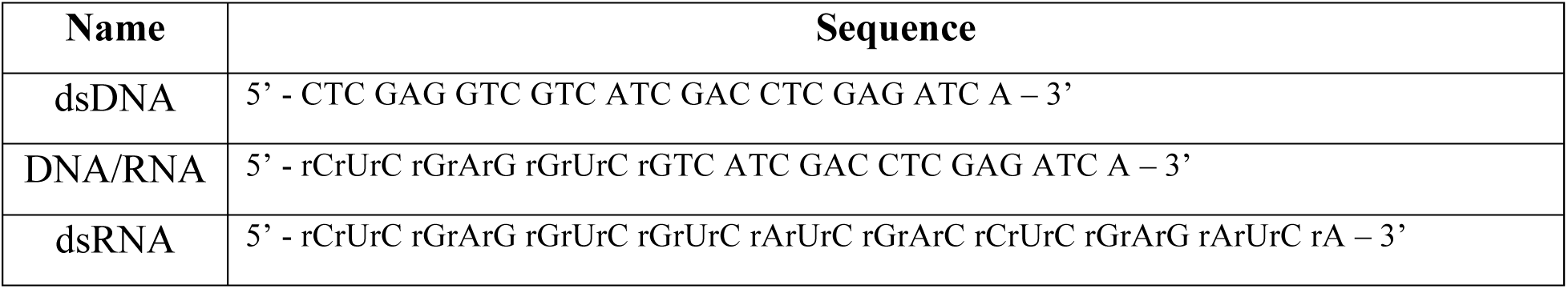

Calculated k_D_ from fitting to hyperbolic equation (Bound = (v*[E])/(k_D_+[E])), where “[E]” is the concentration of the enzyme, and “v” and “k_D_” are solved by linear regression.

### Bioinformatic analysis

#### Protein identification

Prior to protein identification, human and mouse proteomes were downloaded from UniProt website (Apweiler et al., 2004). Raw formatted mass spectrometry files were first converted to mzXML file format using MSConvert (http://proteowizard.sourceforge.net/tools.shtml) and then processed using MSGF+ (Kim et al., 2017), X! TANDEM (Craig and Beavis, 2004) and Comet (Lingner et al., 2011) peptide search engines with default settings. MSBlender (Kwon et al., 2011) was used for integration of peptide identifications and subsequent mapping to protein identifications. A false discovery rate of 1% was used for peptide identification. Protein elution profiles were assembled using unique peptide spectral matches for each protein across all fractions collected.

#### DIF-FRAC score and P-value significance calculation

In order to determine the significance of a protein’s sensitivity to RNase A treatment, we compare the protein’s control elution profile to its RNase A treated elution profile as schematized in Figures 2A and S3A. Specifically, we first calculate the L1-norm of the two elution profiles (equation 1).

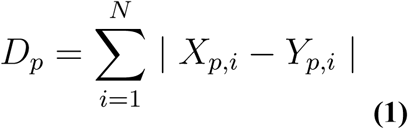

Where *N* represents the total number of fractions collected and *p* represents an individual protein. *X* and *Y* represent abundance matrices of control and experiment (RNase A treated) respectively. We next normalize *D_p_* by the total abundance seen for protein *p* in both the control and experiment (equation 2).

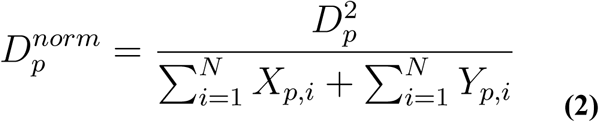

We observed *D^norm^* is biased by high abundance proteins and we therefore evaluate significance of a protein’s sensitivity to RNase A treatment by comparing to a background of proteins with similar abundance. Specifically, we create a distribution of *D*^*norm*^ from proteins in a window surrounding protein *p* and have not been annotated as RNA-associated proteins in the literature (equation 3). See Figure S3A for schematic.

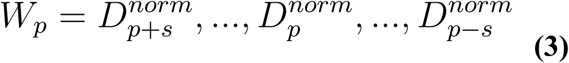

where *s* is a window size of 100 and unannotated RNA-associated proteins are in order of abundance.

We posit that the proteins in distribution, *W_p_*, is a mixture of unannotated RNA-associated proteins as well as non-RNA-associated proteins. In order to evaluate significance of a protein’s *D^norm^* being greater than what is expected by non-RNA binders, we model the distribution *W_p_* using a two component gaussian mixture model (GMM). To ensure an accurate model fit we evaluate our GMM fit using three criteria (equation 4). First, we calculate the Baysian Information Criterion (BIC) for both the two component GMM and a one component GMM and ensure the two component GMM has a lower BIC (equation 4a). Second, we ensure the component with the lowest mean *µ* (i.e. non-RNA-associated component) has the largest weight (equation 4b). Finally, we ensure the largest component weight is greater than a given weight threshold *t_weight_* (equation 4c). *t_weight_* can be estimated by the expected fraction of non-RNA binders in the proteome. In practice we set *t_weight_* to be between 0.6 and 0.75.

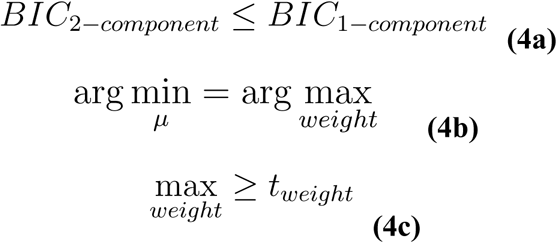

If all three criteria are passed the lowest mean component of the two component GMM is used, otherwise the one component is used (equation 5).

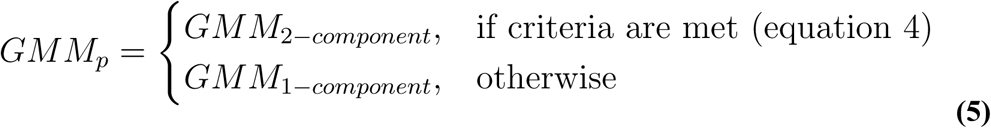

We next calculate the Z-score of protein *p*’s *D^norm^* score relative to the non-RNA-associated component of GMM_p_ (equation 6).

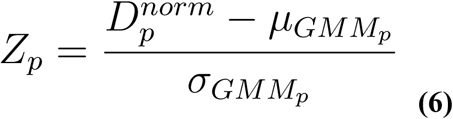

where *µ_GMMp_* and *σ_GMMp_* are the mean and standard deviation of the first component of GMM_p_ respectively.

Finally, we calculate a P-value of *Z_p_* using the normal distribution survival function and then false discovery correct P-values across all proteins using the Benjamini/Hochberg correction (Benjamini and Hochberg, 1995). RNA-associated proteins were considered significant at a 0.05 FDR corrected P-value.

#### RNA-associated annotations, overlap comparisons and score performance analysis

Low throughput RNA binding annotations were defined as proteins with Gene Ontology (Ashburner et al., 2000) “RNA binding” annotations limited to those with evidence codes: EXP, IDA, IPI, IMP, IGI, IEP, TAS, NAS, or IC. In addition, proteins with “ribonucleoprotein” in their UniProt keywords were also included. High throughput RNA association annotations were primarily collected from Hentze *et al*. 2018 supplemental table S2 (Hentze et al., 2018). In addition, we gathered more recent high throughput datasets from (Bao et al., 2018; Huang et al., 2018; Queiroz et al., 2019; Trendel et al., 2019).

To estimate the coverage of identified RNA-associated proteins by the DIF-FRAC method independent of cell type and machine setup, the Venn diagrams in Figures 5B and S4L report only proteins with mean abundance > 10, where mean abundance is the average peptide spectral matches identified in the control and RNase A treated HEK 293T cells. To compare directly the RNA-associated proteins identified in the high throughput sets to the DIF-FRAC method, Venn diagrams in Figure S4A-K report all proteins.

To calculate Precision vs Neg Ln P-value plots (Figure 1I and Figure S3C), we first added a pseudocount (+1e-308) to DIF-FRAC P-values and then applied −1*ln(P-value) where ln is the natural log. Precision is defined as TP/AP, where TP (true positives) is defined as proteins annotated as either high throughput or low throughput RNA binding (see above) and a Neg Ln P-value greater than a given value. AP (all predictions) is defined as any protein with a Neg Ln P-value greater than a given value. To calculate Precision vs Recall plots (Figures S3B and S3D), precision is defined above and recall is defined as TP/AKP where TP is true positives and AKP (all known positives) is defined as proteins annotated as either high throughput or low throughput RNA binding.

#### Classification of DIF-FRAC elution profiles

To calculate the amount a protein shifts upon RNase A treatment, we calculate the average fraction a protein is observed weighted by the PSMs observed in each fraction. The difference between the weighted average of the treated and untreated elution profiles provides the total shift amount. A protein’s shift in elution from a high molecular weight to a low molecular weight results in a negative shift value whereas a shift from low molecular weight to high molecular weight corresponds to a positive value.

To calculate the amount a protein’s abundance changes upon RNase A treatment, we calculate the difference of a protein’s total PSMs observed in the untreated and treated samples. We further normalize this value by dividing by the sum of the total PSMs from both samples. This results in a value between 1.0 and −1.0 where a positive value corresponds to an increase in abundance upon RNase A treatment and a negative value corresponds to a decrease in abundance upon RNase A treatment.

#### Assembly of RNP complexes

We define the global set of RNP complexes by first creating a combined non-redundant set of CORUM (Ruepp et al., 2010) and hu.MAP (Drew et al., 2017) complexes (Jaccard coefficient < 1.0). For every complex in this global set we tested if > 50% of the protein subunits were 1) identified as an RNA-associated protein by DIF-FRAC (P-value > 0.05), 2) annotated by high throughput methods or 3) annotated by low throughput methods (see above for description of annotations). RNP Select complexes are defined as complexes whose protein subunits co-elute in the DIF-FRAC control sample (> 0.75 average Pearson correlation coefficient among subunits) and > 50% of subunits have a DIF-FRAC P-value > 0.5.

#### RNA-Seq Analysis

After performing quality control on the sequencing fastq files using FastQC (www.bioinformatics.babraham.ac.uk/projects/fastqc/), 3’ adapter contamination was removed using Cutadapt (v1.10) (Martin, 2011). Alignment of the 8 RNA fraction datasets was then performed with the Hisat2 transcriptome-aware aligner (v2.1.0) (Kim et al., 2015), against a Hisat2 reference index built using GRCh38/hg38 primary assembly genome fasta from Gencode (v27, Ensembl release 90) (Harrow et al., 2012) annotated with the corresponding v27 GTF (General Transfer Format) annotations. The Hisat suite Stringtie program (v1.3.3b) (Pertea et al., 2016) was used to quantify gene-level expression from the alignment files. TPM (Transcripts Per Million), a sequencing-depth-normalized estimate of reads mapping to the gene, was used for further analysis.

#### Data Deposition

Proteomics data and RNA-seq data will be deposited in Pride and Gene Expression Omnibus respectively upon acceptance.

#### Code Repository

Source code is freely available on GitHub: https://github.com/marcottelab/diffrac

---

**Figure S1:**
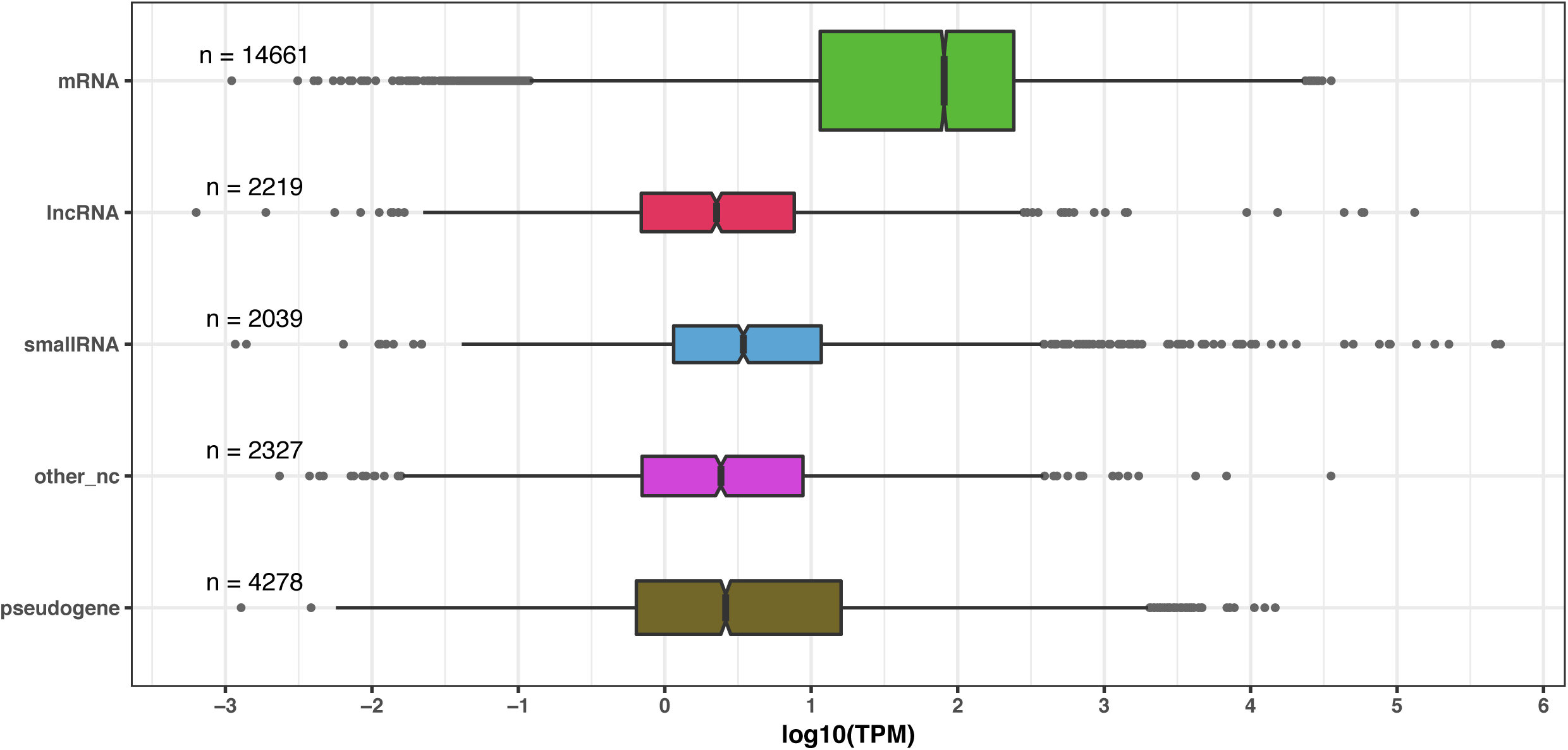
DIF-FRAC accesses a diverse RNA landscape. Box plots show the RNA abundance of mRNA, lncRNA, small RNA, other ncRNA, and pseudogenes in control fractions 16-23 of HEK 293T cell lysate (TPM = Transcripts Per Million). Boxes indicate median (inner joint), first quartile (left) and third quartile (right). Lines indicate 1.5 interquartile range. Dots indicate outliers.

**Figure S2:**
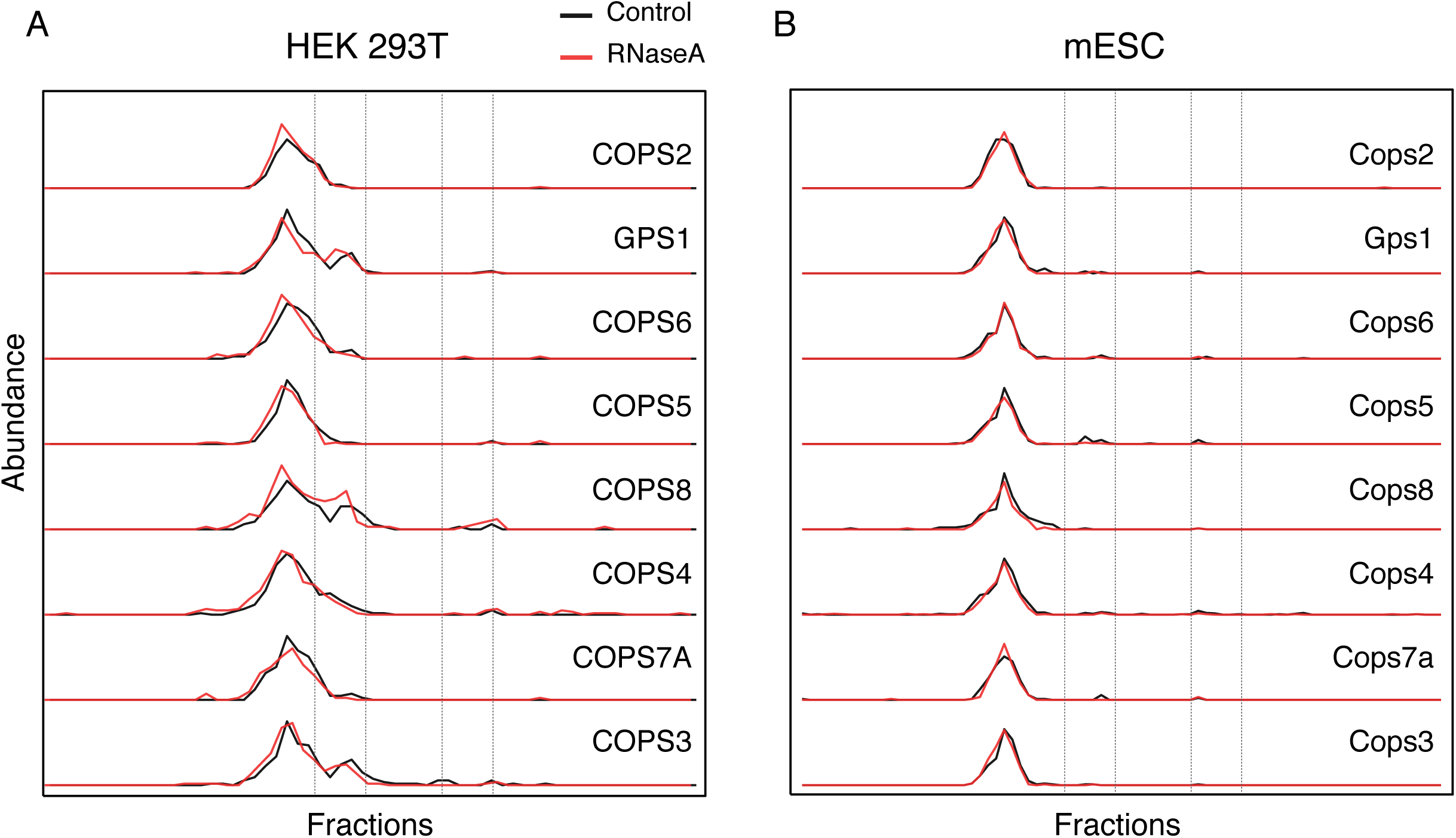
Non-RNA-associated complexes are insensitive to RNase A treatment. DIF-FRAC elution profiles show subunits of the negative control non-RNA-associated COP9 signalosome complex (M_r_ ∼500 kDa (Oron et al., 2002)) in control (black) and RNase A treated (red) for (A) HEK293T lysate, and (B) mESC do not shift upon RNase A treatment. Abundance represents count of unique peptide spectral matches. Vertical dotted lines represent protein standards described in Figure 1.

**Figure S3:**
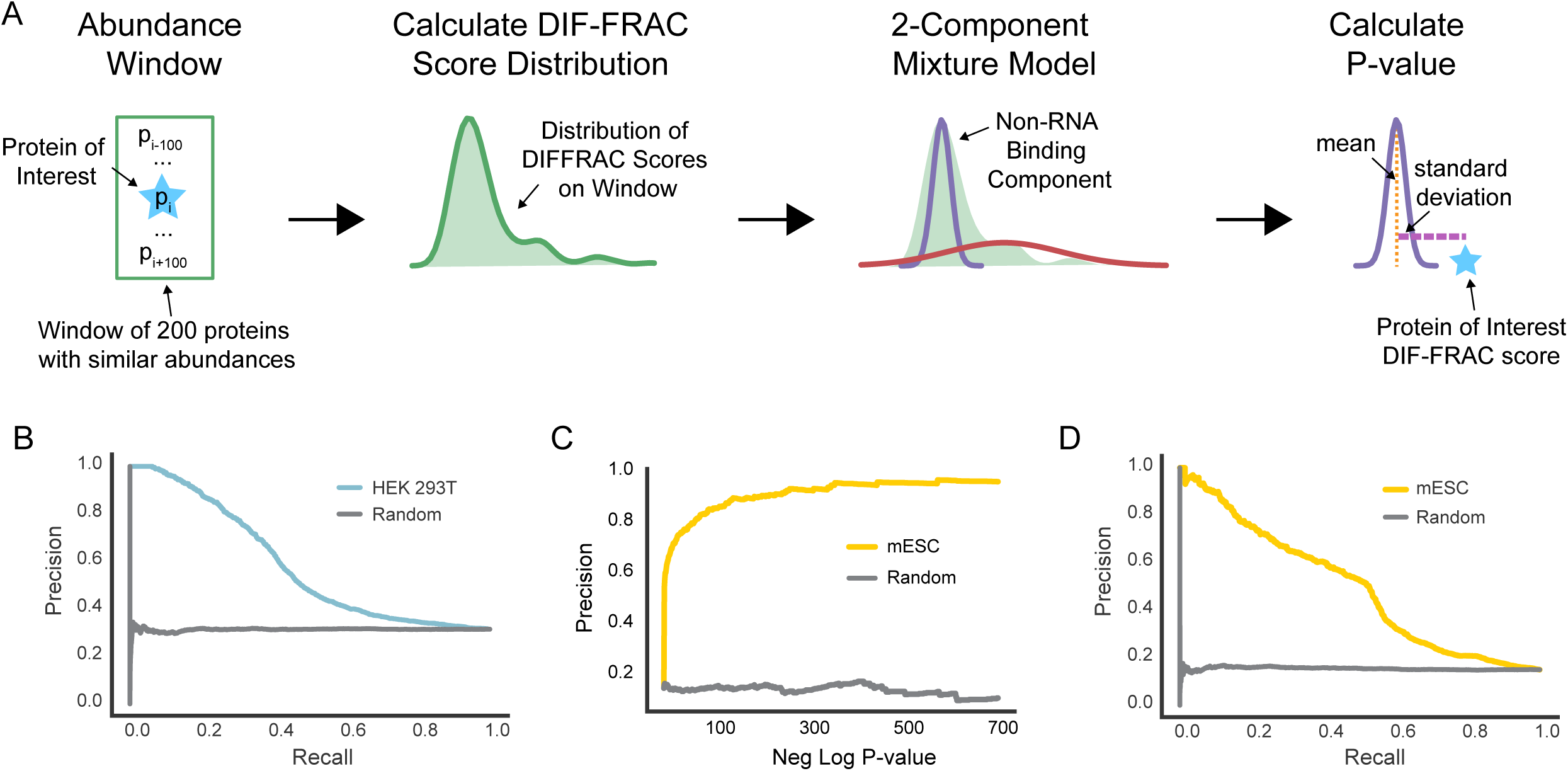
DIF-FRAC Score accurately discriminates between RNA-associated proteins and non-binders. (A) Workflow to calculate abundance corrected P-values for each protein’s DIF-FRAC score. Proteins are ranked according to abundance and a window of +/- 100 proteins is used to calculate a DIF-FRAC score distribution. A two-component Gaussian mixture model is then used to identify the non-RNA binding component in the distribution. Finally, the DIF-FRAC score of the protein of interest is compared to the non-RNA binding distribution component to test the null hypothesis and a P-value is calculated. (B) Precision recall analysis shows the DIF-FRAC Score recalls a substantial number of known RNA-associated proteins in HEK 293T cells. (C) High DIF-FRAC P-values have high precision in recovering known RNA-associated proteins in mouse embryonic stem cells. (D) Precision recall analysis shows the DIF-FRAC Score recalls a substantial number of known RNA-associated proteins in mESC.

**Figure S4:**
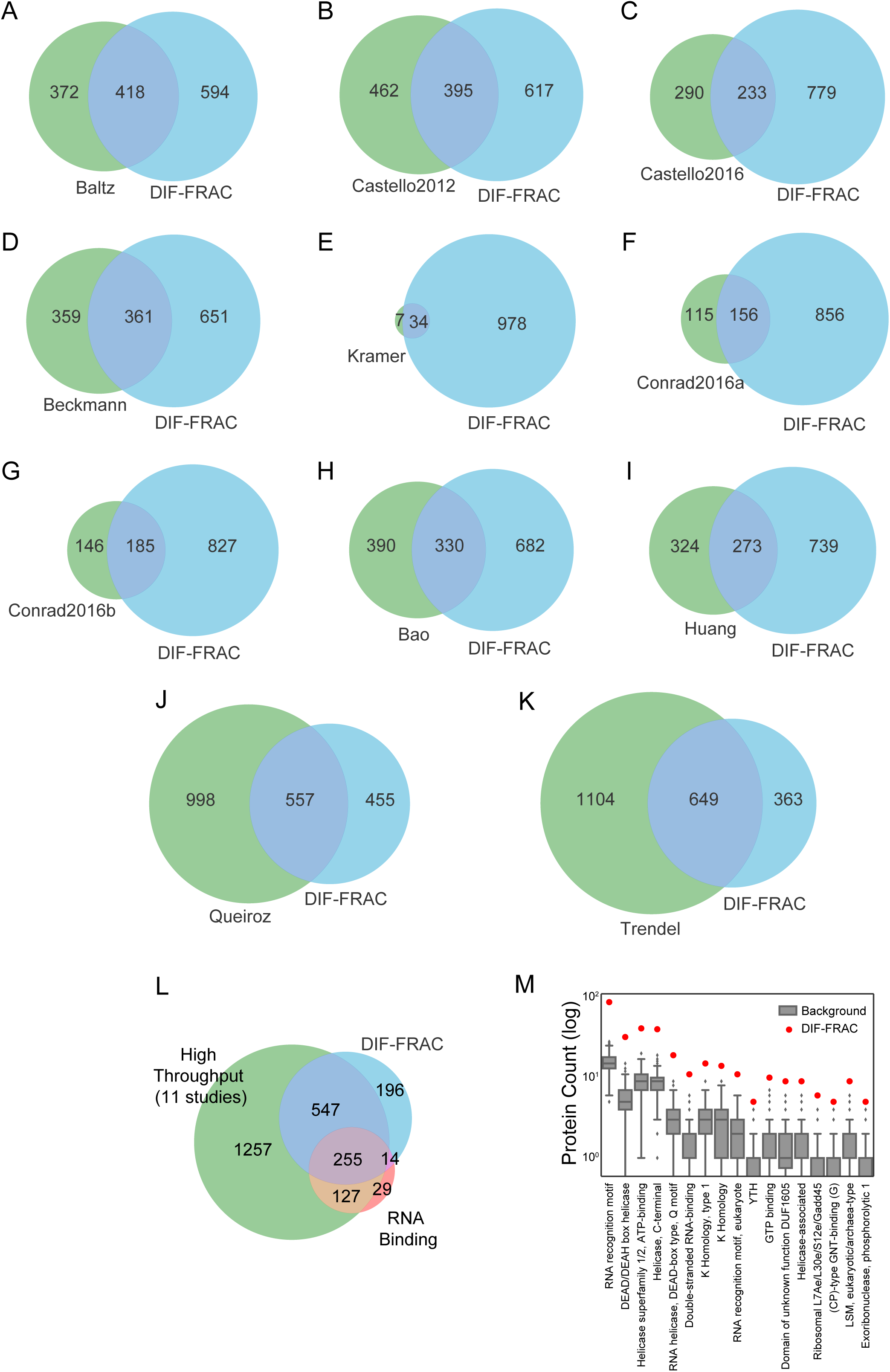
DIF-FRAC RNA-associated proteins show substantial overlap with other high-throughput studies. (A-K) Venn diagrams show overlap of DIF-FRAC RNA-associated proteins from HEK 293T cells (blue) with 11 high-throughput RNA association studies (green) (Bao et al., 2018; Hentze et al., 2018; Huang et al., 2018; Queiroz et al., 2019; Trendel et al., 2019). (L) Venn diagram shows overlap of DIF-FRAC RNA-associated proteins from HEK 293T cells (blue), annotated RNA Binding proteins (red), and combined set of high throughput RNA association studies (green). (M) Enrichment of RNA binding structural motifs in DIF-FRAC-identified RNA-associated proteins from HEK 293T cells.

**Figure S5:**
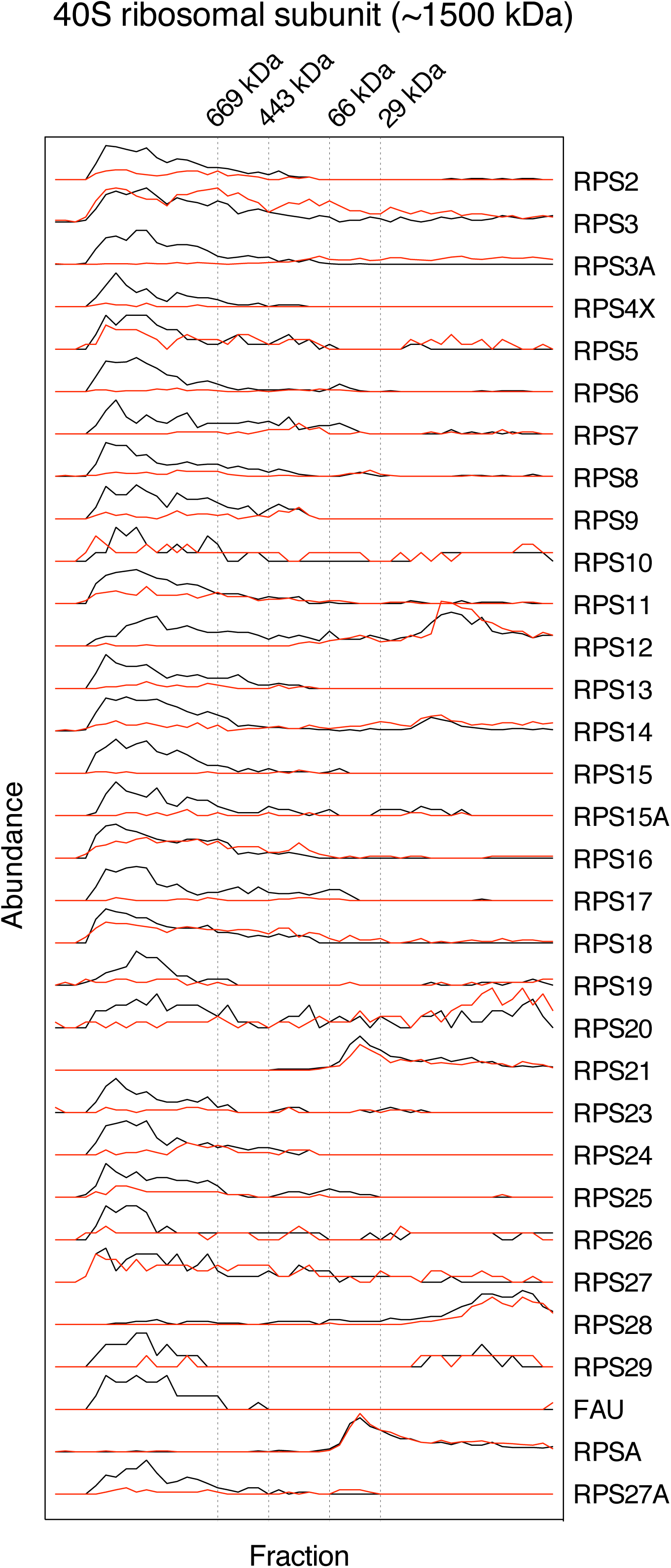
DIF-FRAC classifies the 40S ribosomal subunit as a structural RNP. Elution profiles of the 40S ribosomal subunits demonstrate it is destabilized upon RNA degradation (‘structural’ RNP complex).

**Figure S6:**
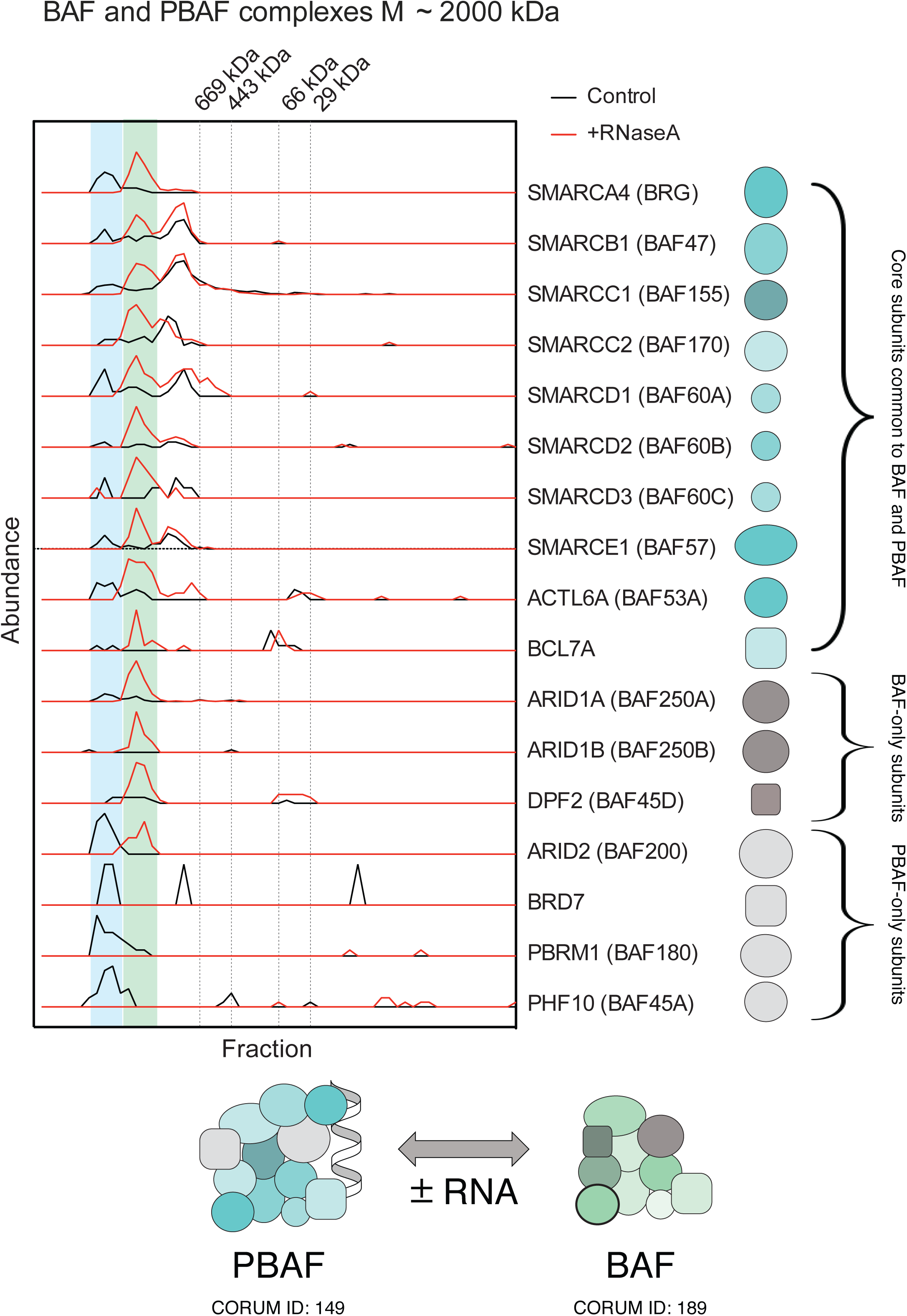
DIF-FRAC classifies human BAF and PBAF complexes as compositional RNPs. Elution profiles of annotated human PBAF (blue shading) and BAF (green shading) complexes demonstrate core subunits common to both complexes coelute in both control and RNase A treated samples, but at different molecular weights. PBAF-only subunits (light grey) coelute with core subunits only in the control sample, while BAF-only subunits (dark grey) coelute with the core subunits as a lower molecular weight complex only when RNA is degraded. Together, these elution profiles suggest that PBAF is an RNP complex, but the BAF complex does not associate with RNA.

**Figure S7:**
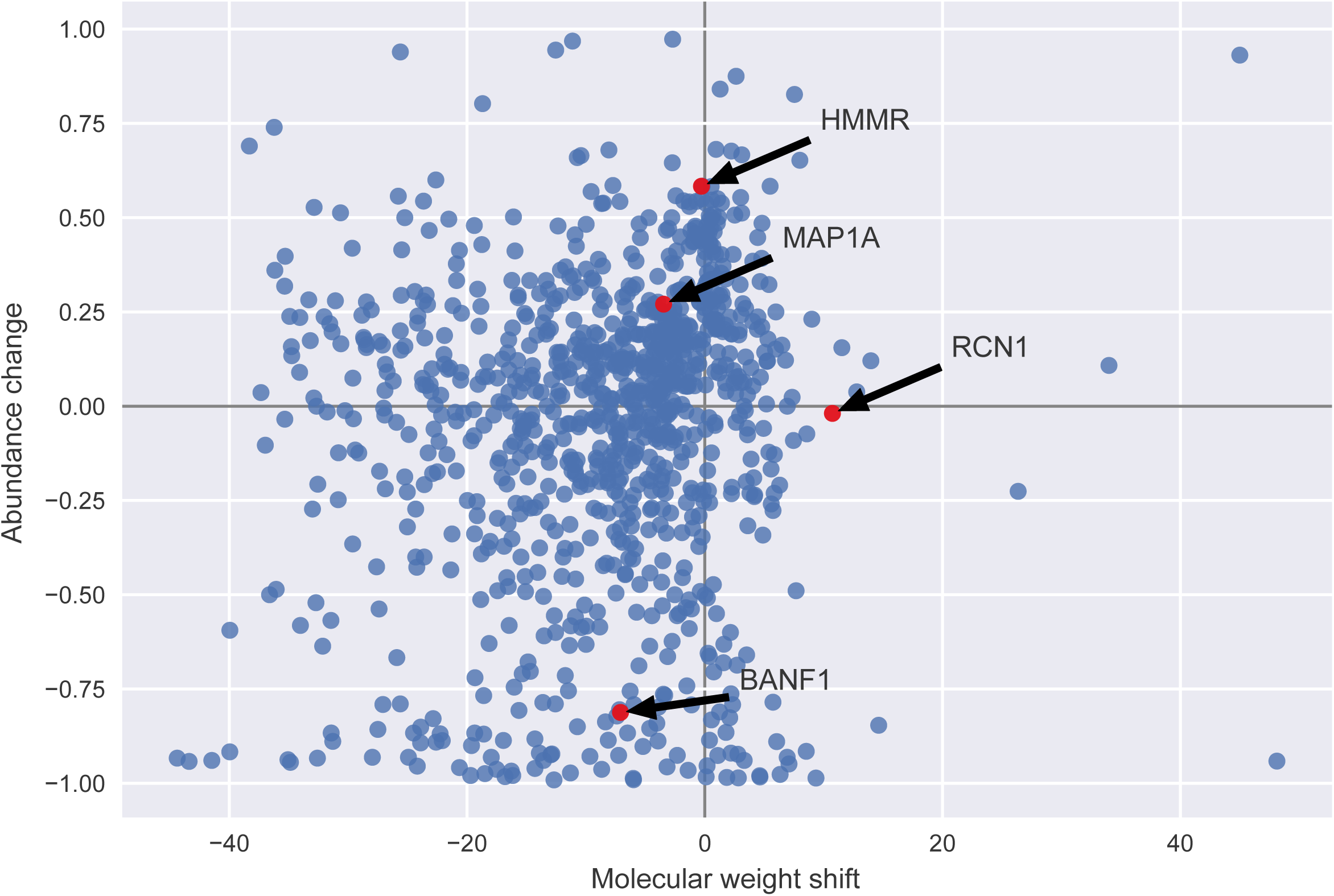
Analysis of DIF-FRAC shift types of RNA-associated proteins. Upon RNase A treatment we observe different types of changes to protein elution profiles (see Figure 1H and Figure 4). Each point in the graph represents one RNA-associated protein. Molecular weight shift is the weighted average difference between control and RNase A treated profiles, where a negative value (left side of graph) represents lower molecular weight elution upon treatment and positive value (right side of graph) represents gain in molecular weight (see Methods for calculation). Abundance change is the normalized change in observed abundance upon RNase A treatment. A positive value (top of graph) represents gain in solubility and a negative value (bottom of graph) represents loss in solubility. Examples from Figure 4 are annotated.

**Figure S8:**
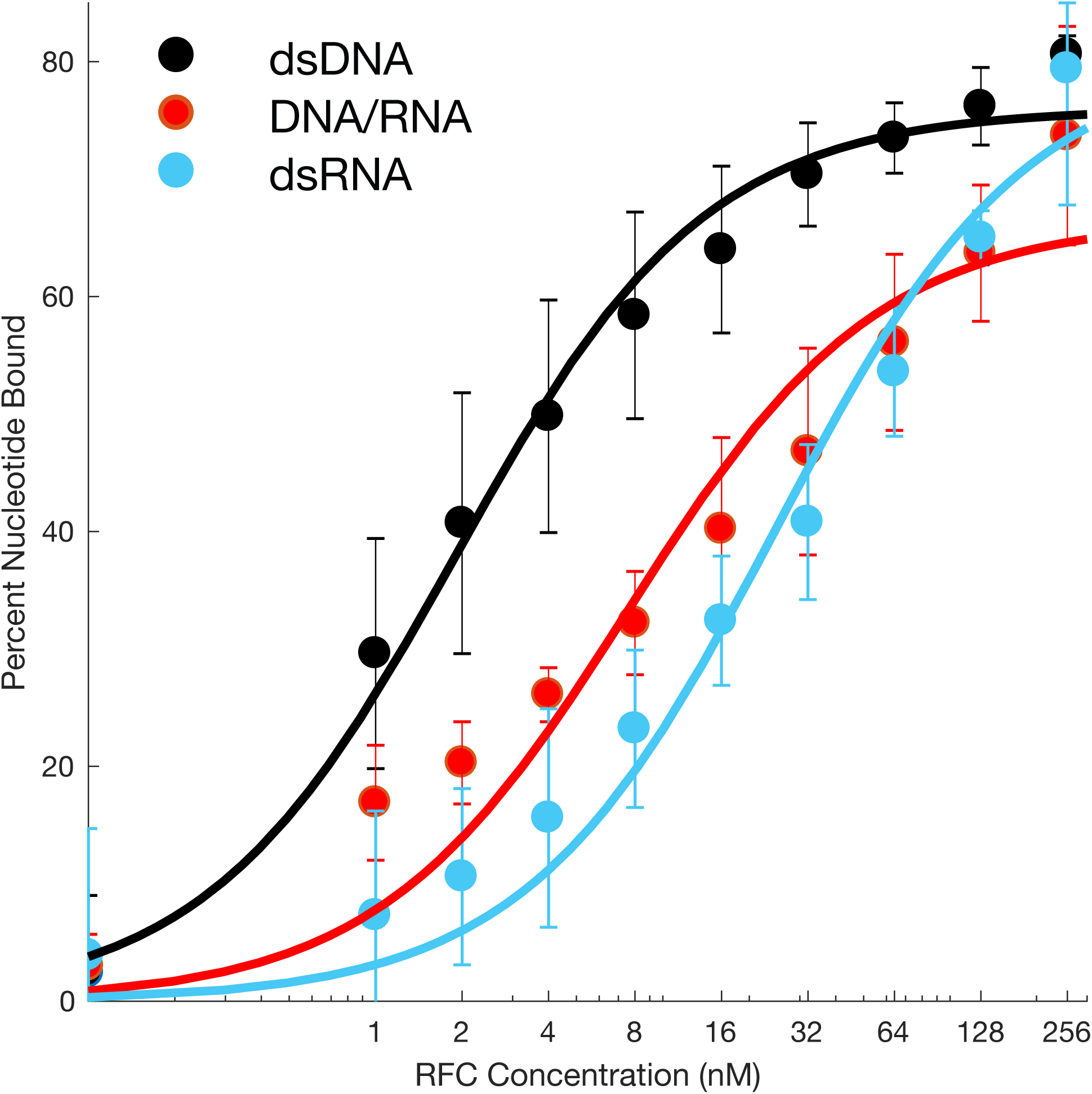
Affinity of nucleic acid for the *S. cerevisiae* RFC complex. Binding curves from electromorphic mobility shift assays (EMSA) of various concentrations of purified *S. cerevisiae* RFC mixed with 1 nM ^32^P-labeled oligonucleotides. Data was fit to a hyperbolic equation (solid line). The calculated k_D_ + 95% CI is 1.9 ± 0.5 nM for dsDNA (black), 7.5 ± 4.9 nM for DNA/RNA hybrid (red), and 25 ± 11 nM for dsRNA (blue). Error bars denote standard deviation.

