## Supplemental Table 2 for "Systematic discovery of endogenous human ribonucleoprotein complexes"

### Table S2: Top 15 unannotated RNA-associated proteins identified by DIF-FRAC

| | Gene Name | Protein | Function | Soluble without RNA? $ | Disease links # | DIF-FRAC score/ p-value (5 % FDR) | DIF-FRAC plot |
|---|---|---|---|---|---|---|---|
| 1. | BANF1 | Barrier-to-autointegration factor | Chromatin organization | No | Progeria syndrome | 6.17E-45 | 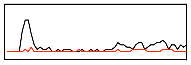 |
| 2. | RCN1 | Reticulocalbin-1 | Calcium binding<br>Secretory pathway<br>Stress Response | No | Amyloid formation<br>Hepatocellular carcinoma | 4.97E-41 | 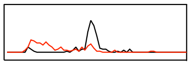 |
| 3. | MAP1A | Microtubule-associated protein 1A | Microtubule assembly<br>Structural protein | Yes | Hearing loss | 1.38E-39 | 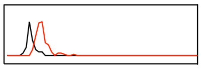 |
| 4. | NOMO3 | Nodal modulator 3 | Carbohydrate binding | Yes | N/A | 2.29E-32 | 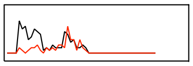 |
| 5. | ACTA2 | Actin, aortic smooth muscle | Muscle protein | No | Vascular diseases | 7.11E-31 | 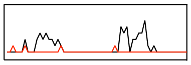 |
| 6. | RSBN1L | Round spermatid basic protein 1-like protein | N/A | No | N/A | 1.45E-27 | 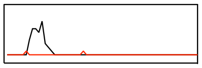 |
| 7. | NIPSNAP1 | Protein NipSnap homolog 1 | Neurotransmitter binding | Yes | N/A | 1.92E-24 | 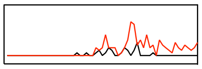 |
| 8. | CLSPN | Claspin | DNA binding<br>DNA replication | No | N/A | 2.79E-22 | 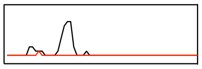 |
| 9. | U2AF1L5 | Splicing factor U2AF 35 kDa subunit-like protein | RNA binding (by similarity) | No | N/A | 3.76E-22 | 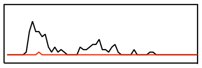 |
| 10. | MORF4L2 | Mortality factor 4-like protein 2 | Chromatin regulator | Yes | N/A | 9.82E-22 | 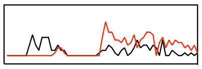 |
| 11. | HMMR | Hyaluronan mediated motility receptor | Hyaluronic acid binding | Yes | Breast cancer | 2.04E-17 | 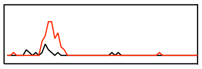 |
| 12. | PTGES2 | Prostaglandin E synthase 2 | Isomerase | Yes | Type 2 diabetes | 2.52E-17 | 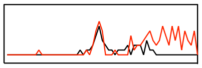 |
| 13. | WNK2 | Serine/threonine-protein kinase WNK2 | Serine/threonine-protein kinase | Yes | N/A | 4.14E-17 | 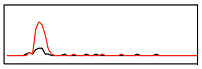 |
| 14. | MARK3 | MAP/microtubule affinity-regulating kinase 3 | Serine/threonine-protein kinase | Yes | Pancreas carcinogenesis | 1.40E-16 | 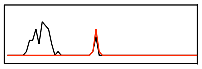 |
| 15. | ALOX5 | Arachidonate 5-lipoxygenase | Leukotriene biosynthesis | Yes | Asthma | 1.21E-15 | 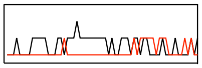 |

$ Insolubility in the absence of RNA is inferred by an increase in elution volume/ molecular weight of the protein upon RNA digestion, or a complete disappearance of signal. This is consistent with the RNA-associated protein being solubilized by RNA, as suggested by Maharana et al. (Maharana et al., 2018)

### Annotations from UniProt (The UniProt, 2017) and/ or OMIM (https://omim.org/)
